# Numerical methods for the detection of phase defect structures in excitable media

**DOI:** 10.1101/2021.12.16.473086

**Authors:** Desmond Kabus, Louise Arno, Lore Leenknegt, Alexander V. Panfilov, Hans Dierckx

**Affiliations:** Department of Mathematics, KU Leuven Campus Kortrijk (KULAK), Kortrijk, Belgium; Laboratory of Experimental Cardiology, Leiden University Medical Center (LUMC), Leiden, The Netherlands; iSi Health, KU Leuven Institute of Physics-based Modeling for In Silico Health, Leuven, Belgium; Department of Physics and Astronomy, Ghent University, Ghent, Belgium; Laboratory of Computational Biology and Medicine, Ural Federal University, Ekaterinburg, Russia; World-Class Research Center “Digital biodesign and personalized healthcare”, Sechenov University, Moscow, Russia

## Abstract

Electrical waves that rotate in the heart organize dangerous cardiac arrhythmias. Finding the region around which such rotation occurs is one of the most important practical questions for arrhythmia management. For many years, the main method for finding such regions was so-called phase mapping, in which a continuous phase was assigned to points in the heart based on their excitation status and defining the rotation region as a point of phase singularity. Recent analysis, however, showed that in many rotation regimes there exist phase discontinuities and the region of rotation must be defined not as a point of phase singularity, but as a phase defect line. In this paper, we use this novel methodology and perform a comparative study of three different phase definitions applied to *in silico* data and to experimental data obtained from optical voltage mapping experiments on monolayers of human atrial myocytes. We introduce new phase defect detection algorithms and compare them with those that appeared in literature already. We find that the phase definition is more important than the algorithm to identify sudden spatial phase variations. Sharp phase defect lines can be obtained from a phase derived from local activation times observed during one cycle of arrhythmia. Alternatively, similar quality can be obtained from a reparameterization of the classical phase obtained from observation of a single timeframe of transmembrane potential. We found that the phase defect line length was (35.9 *±* 6.2) mm in the Fenton-Karma model and (4.01 ± 0.55) mm in cardiac human atrial myocyte monolayers. As local activation times are obtained during standard clinical cardiac mapping, the methods are also suitable to be applied to clinical datasets. All studied methods are publicly available and can be downloaded from an institutional web-server.

## 1 Introduction

The heart is a self-organizing dynamical system for which the mechanical contraction is regulated by waves of electrical activation travelling through the cardiac wall. During cardiac arrhythmia complex electrical patterns emerge that often result into a rotating pattern, either circling around an obstacle or around its own wave back [1, 2]. These vortices are also known as rotors, or spiral waves in two dimensions (2D), or scroll waves in three dimensions (3D).

After the experimental observation of such structures in animal hearts during ventricular tachycardia [3], it was conjectured that rotors can sustain several heart rhythm disorders. However, the precise dynamics of rotors, the structure of the rotor core and the most efficient manner to remove them from the heart remain incompletely understood.

A quantitative description of spiral wave motion requires localizing it in space. In first approximation, the region around which the rotor revolves is called the spiral wave core. The location of a spiral wave at a given time can be further narrowed down to a single point, usually called the spiral wave tip. Different methods exist to define the tip, e.g. as the point where wave front and wave back merge [4], as the point on a line of constant voltage that does not change instantaneously, as an intersection between two isolines of different variables [5] or as a point singularity of the activation phase [2, 6].

By following the tip position of a single spiral wave over time, a disc-like or star-like shape emerges, known as the spiral wave core. Different types of cores have been observed [7], and the non-circular cores are referred to as meandering cores. Among the meandering cores, simulations of detailed ionic models for cardiac tissue typically show so-called linear cores, as depicted in Fig 1.

**Fig 1.**
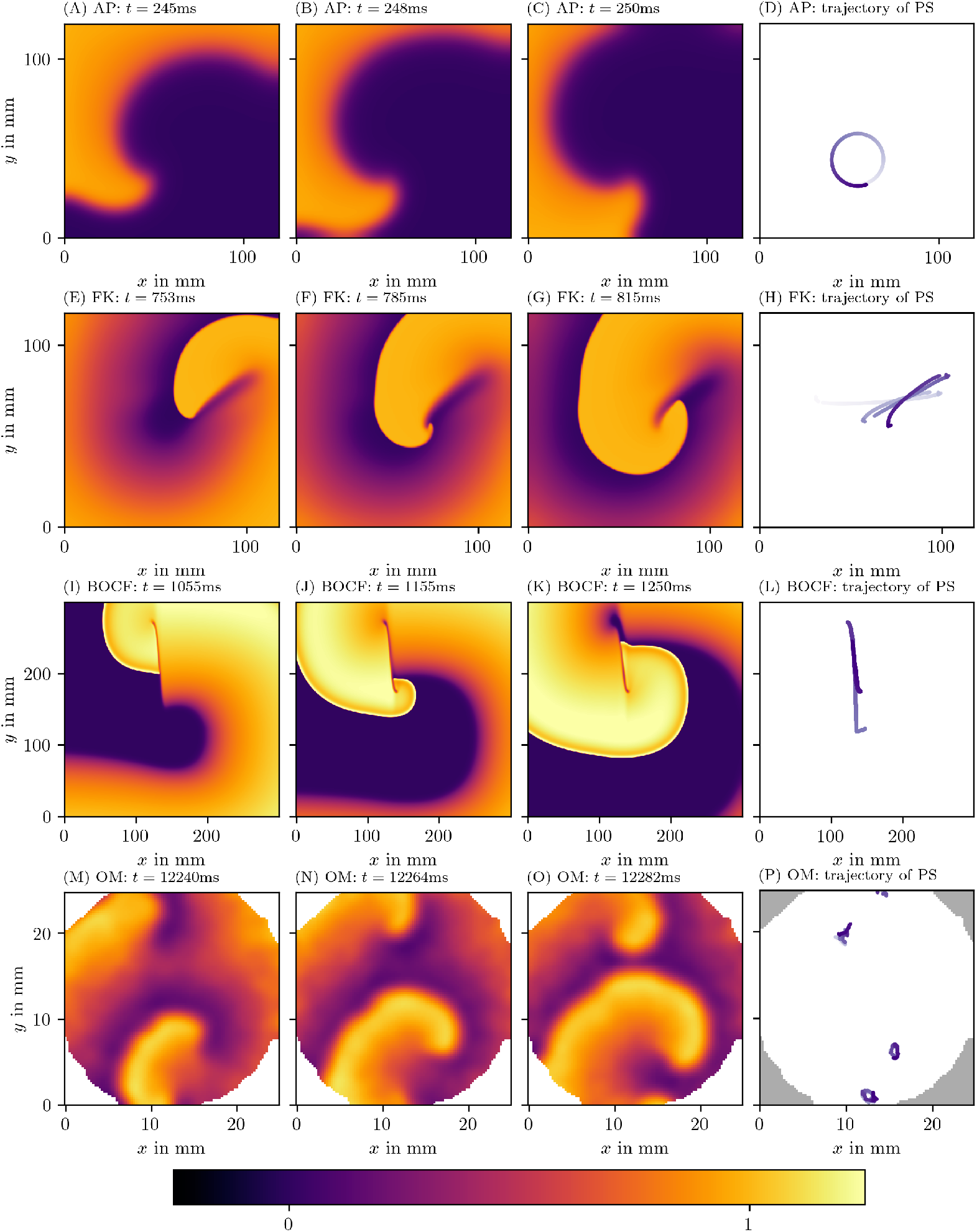
Qualitatively different tip trajectories of spiral waves in modelled cardiac tissue. The first row displays the Aliev-Panfilov (AP) model [8], the second row the Fenton-Karma (FK) model [5] with modified Luo-Rudy I (MLR-I) parameters, the third row Bueno-Orovio-Cherry-Fenton (BOCF) [9], and the last row displays optical voltage mapping data derived from monolayers of conditionally immortalized human atrial myocytes (hiAMs) [10]. In section 3, we outline in more detail how these data sets have been obtained. The first 3 columns show different snapshots in time and are colored according to local transmembrane voltage. Corresponding spiral wave tip trajectories are shown in the last column. Darker coloring of the trajectory corresponds to later points in time.

The linear-core regime arises in systems with long action potential duration (APD): If the tip is next to a region of refractory tissue, it will follow this interface until meeting a point where the tissue has recovered. As a result, the tip moves along an almost straight line, interleaved with turning points. These dynamics have not only been observed in simulations, but were also reported in experiments [11, 12] in the form of a line of conduction block. For this reason, it is important to elucidate the spatial distribution of the core of the rotor.

In 3D, a spiral wave or rotor becomes a structure called a scroll wave. Within the scroll wave, the collection of spiral wave tips forms a filament curve [13]. In numerical simulations with linear cores and a few experimental observations [11], the straight segment in the linear core extends to a ribbon-like filament [11]. However, these spatially extended filaments have not been substantially included in theory development, as was done for circular-core filaments [14].

Recent works [15, 16] have proposed to treat linear cores fundamentally different from circular cores. Specifically, classical phase analysis [2, 6] of cardiac activation patterns assumes that there is a phase singularity (PS) located near the spiral wave tip. However, when the wave front reaches a part of tissue that is for instance not fully recovered yet, a so-called conduction block will form. On both sides of this conduction block line, there is a different phase, such that the authors of [15, 16] argue that the phase representation resembles more a phase defect line (PDL), i.e. a line where the phase changes abruptly from one side to the other. In other words, we go from a phase singularity (PS) to a branch cut known in complex analysis. Making the distinction between phase defects (PDs) and point singularities could have potential clinical use, as current analysis methods in simulation, experiment and of clinical data are only aiming to localize point singularities of phase.

The aim of this paper is to provide and compare several methods to numerically calculate the phase and the corresponding PDs. We include a quantitative analysis of the results and the performance of the algorithms.

A summary of our workflow is presented in Fig 2: An image (e.g. of the transmembrane potential *u*) is in the first step converted into a phase *ϕ*. Where jumps in phase are detected, the PD density *ρ* (see below) will be much larger than zero. If desired, the field 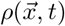 can be further processed to yield localized PDLs.

**Fig 2.**
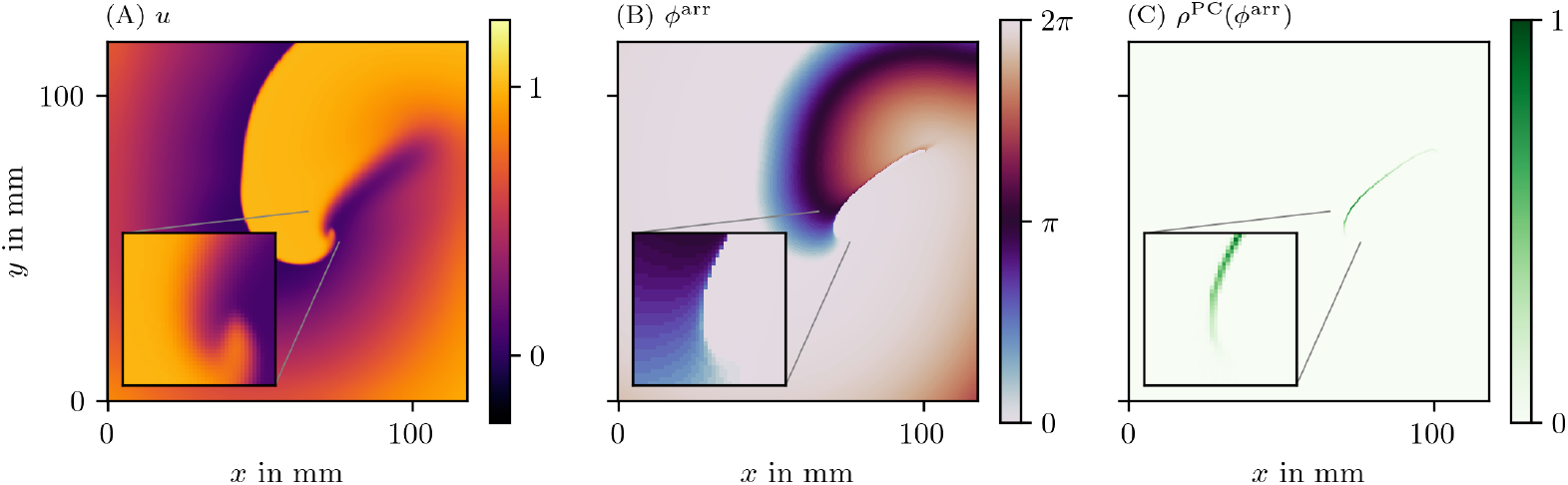
An example of the steps done in the process of constructing the PD density *ρ*. (A) We start from the first phase state variable *u*, often the normalized and hence unitless transmembrane potential *V*. (B) Next, from this *u* and possibly other state variables, the phase, here *ϕ*^LAT^, is calculated. How to compute it will be introduced below in Eq (3). (C) Finally from the phase, the PD density is produced based on one of the PDL detection algorithms; here, the phase coherence (PC) method was used, which will be introduced below in section 3.3.3. These quantities are plotted over a 2D square domain of myocardial tissue in physical space. Color is used to represent the values of the quantities mentioned in the top left corner. The same coloring will be used throughout this manuscript.

This manuscript is organized as follows: First, we briefly review the concepts of phase, PSs and PDs (section 2). In the methods section 3, we outline our simulation methods and present several methods to trace PDs in excitation patterns. Results of these methods and their performance are presented in section 4. We conclude this paper with a discussion and outlook (sections 5 and 6).

## 2 Theoretical background

### 2.1 The concept of phase

As remarked long ago by Winfree [17], many biological processes take values within a cycle rather than on the line of real numbers. Cardiac excitation is such an example, since during an action potential, the cell membranes depolarize and repolarize in normal circumstances along a predefined sequence, tracing out a closed loop in state space. To keep track of the relative state of cells along this cycle, the concept of phase can be used. In addition to the classical phase definition, called activation phase *ϕ*^act^ below, an alternative phase based on local activation times (LAT) was defined by Arno *et al*. [16]. In this paper, a third phase *ϕ*^skew^ will be defined below as an approximation of *ϕ*^act^ when LAT are not available (Eq (7)).

Next, we will briefly review the previously defined phases *ϕ*^act^ and *ϕ*^LAT^.

The activation phase *ϕ*^act^ is the phase as seen in a space spanned by two observables 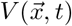 and 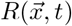 in the system [2, 6, 13]. We henceforth assume that *V* is representing the activation or depolarization of the medium, i.e. in cardiac context, we take *V* to be the normalized transmembrane potential.

Even if there is only one variable *V* observed, its time-delayed version [3], time derivative [5], or Hilbert transform [6] can be used as a linearly independent variable *R*. In numerical simulations, all state variables of the system can be observed, and any pair can be chosen as (*V, R*). Then, one usually defines the activation phase as the polar angle of a state in the (*V, R*)-plane, relative to a reference point (*V*_*_, *R*_*_) that lies within the cycle:

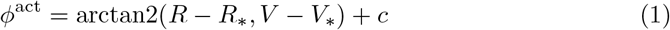

Here, the polar angle is returned by the two-argument inverse tangent: arctan2(*y, x*) = arctan (*y/x*) if *x* ≥ 0 and arctan(*y/x*) + *π* mod 2*π* if *x <* 0. A constant *c* can furthermore be added to make *ϕ*^act^ = 0 correspond to the resting state. The left column of Fig 3 visualizes *ϕ*^act^ for the four data sets introduced in Fig 1.

**Fig 3.**
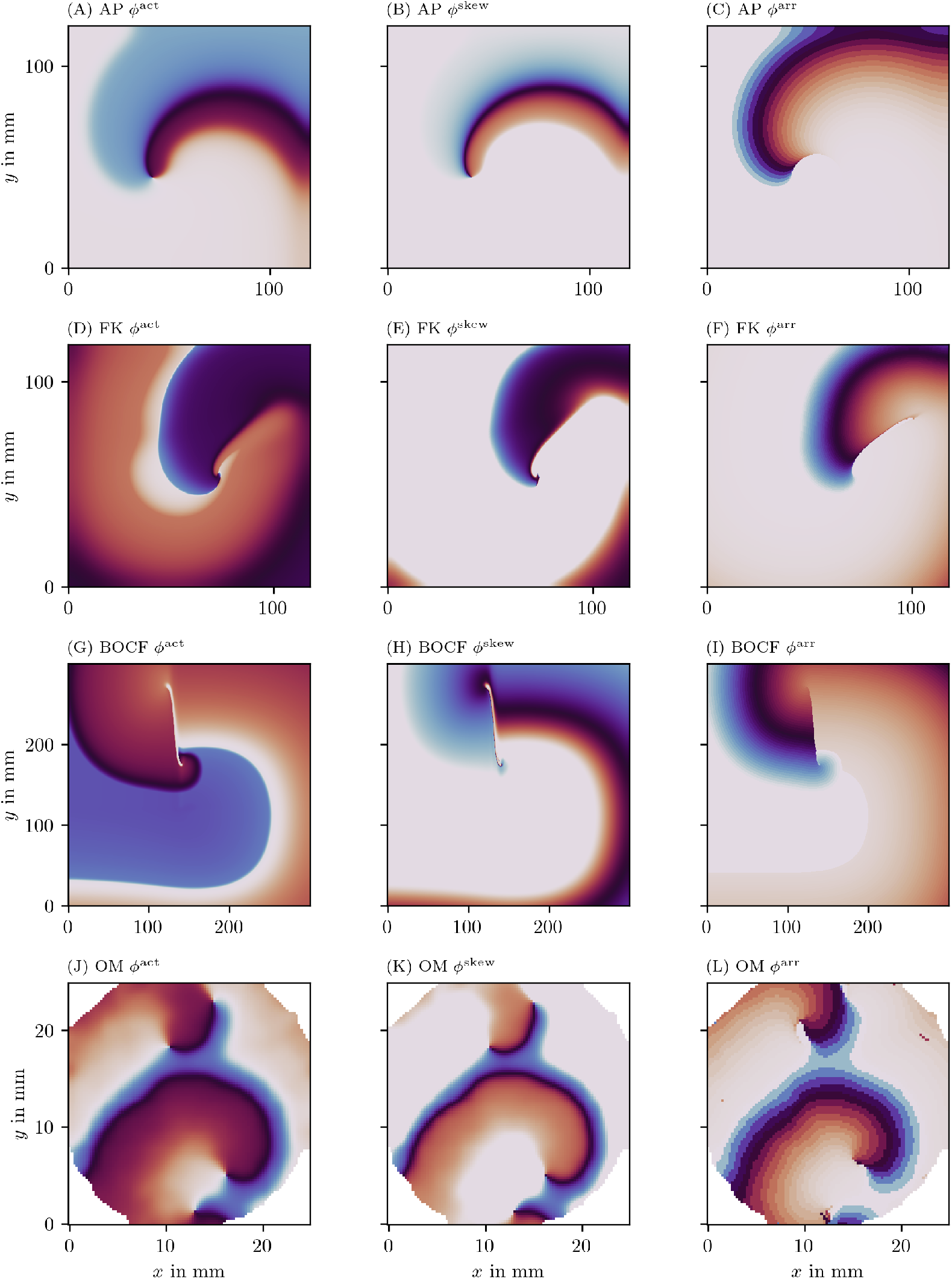
Illustration of the three different phases for one frame of four data sets. The color code represents the phase in modelled myocardial tissue in a 2D square domain in the same scale as in Fig 2B.

We recently proposed a second definition of phase that is based on the local activation time (LAT) of tissue [18]. The LAT, which is commonly used clinically, is defined as the time *t*^*arrival*^ when the tissue locally depolarizes, i.e. when the transmembrane voltage *V* at that point exceeds a value *V*_*\**_.

In noisy circumstances (see Fig 12 below), care is taken to estimate LAT in a robust manner by using an alternative definition of LAT with two thresholds: It has another condition that needs to be met for LAT to be updated. This condition is that *V* must first decrease below a second threshold 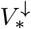 before an increase above the first threshold *V* is considered as depolarization at the wave front and therefore as the trigger of an update of LAT. Unless noted otherwise, we use the first definition of LAT with just one threshold.

The LAT relative to the current time is the elapsed time:

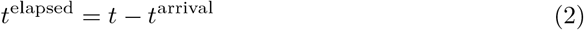

In other words, elapsed time is the time since the start of the last local activation.

The *LAT phase ϕ*^LAT^ is just a mapping of *t*^elapsed^ onto the interval [0, 2*π*) by applying a sigmoidal function:

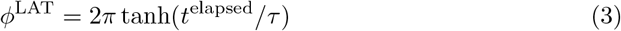

where *τ* is the characteristic time of the cyclic process. Here, we take *τ* equal to half of the typical APD in the medium. Note that another sigmoid function instead of tanh could also have been chosen in Eq (3). The right column of Fig 3 visualizes *ϕ*^LAT^ for the same four data sets.

An asset of *ϕ*^LAT^ is that the curves of equal *ϕ*^LAT^ are precisely the isochrones that cardiologists work with during endocardial catheter mapping. Also, if *τ* is chosen appropriately, repolarized (recovered) tissue will have *ϕ*^LAT^ ≈ 2*π*, such that this phase is not changing abruptly at the wave front and wave back, in contrast to *ϕ*^act^. A disadvantage of *ϕ*^LAT^ is that it requires a dense temporal sampling (either in simulation or experiment), since otherwise staircase artifacts emerge. As a last remark, care should be taken when initializing *ϕ*^LAT^ at the start of the observation window: If the previous LAT are unknown, so is *ϕ*^LAT^ until a propagating wave has swept through the medium.

### 2.2 Phase singularities (PSs) and phase defects (PDs)

The analysis of spatial distributions of phase makes use of several concepts of complex numbers and complex analysis [19], such as contours, PSs and branch cuts. We now briefly review these concepts in the context of cardiac excitation.

The detection of rotor cores from a spatial map of phase can be performed by calculating the total phase difference along a closed loop (contour) *𝒞* in the medium, usually taken on the cardiac surface:

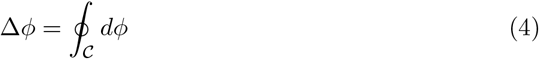

Since the first and last point have the same phase, the resulting phase difference Δ*ϕ* will return an integer multiple of 2*π*. Hence, one defines the topological charge circumscribed by the contour as:

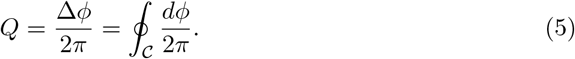

Now, in the classical theory [3], one assumes that the phase function 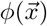 at a given time *t* is continuous nearly everywhere, except in a few points where the phase is undefined. When using *ϕ*^act^, it can be seen that these points will correspond to *V* = *V*_*\**_, *R* = *R*_*\**_. In the immediate vicinity of such points, all phases are present (both in the (*V, R*) plane and in the spatial phase map), hence this point is called a phase singularity (PS). Note that the existence of a PS, where all different phases touch, within the contour region *C* is implied by the *assumption* of the original theory that the phase is a continuous function except at the PS.

In the generalized theory [15, 18], however, the phase is allowed to be discontinuous near a conduction block line, which also happens in the core of a linear-core spiral. Then, the contour 𝒞 cannot be shrunk to surround a single point, without having the contour cross an interface where *ϕ* changes abruptly. We call such transition zones phase defects (PD): phase defect lines (PDLs) in 2D and phase defect surfaces (PDSs) in 3D. The discontinuous behavior of phase is more easily noted with *ϕ*^LAT^ than with *ϕ*^act^, since *ϕ*^act^ also shows strong gradients near the wave front and wave back [18]. As a result, we are convinced that PS detection algorithms that assume a continuous phase distribution are behaving non-robustly near such spatial phase transition, which motivates this work. Here, we will provide PD detection methods that are explicitly discriminating the PD structures, either as a sharp line, or in a probabilistic manner using PD density.

## 3 Methods

### 3.1 Data generation and collection

#### 3.1.1 Numerical methods for pattern generation

The methods for PD detection developed here are designed to operate on excitation patterns, regardless of their generation. However, we here test the methods on numerical simulations in a cardiac monodomain setting. That is, in a rectangular Cartesian grid, we modeled forward evolution of a column matrix of state variables 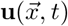 according to a reaction-diffusion equation [20]:

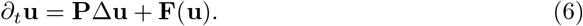

Here, the first component *u*_1_ of **u** equals the normalized transmembrane potential *u*_1_ = *u* = *V*, and **P** = diag(1, 0, …, 0) in order to enable wave propagation by diffusion of *u*. The number of state variables in **u** varies between the different mathematical models of cardiac myocytes, which are encoded in non-linear reaction functions **F**(**u**). To assess the reliability of our methods, we tried several reaction kinetics: Linear cores are known to occur with the Bueno-Orovio-Cherry-Fenton (BOCF) model for human ventricles [9] with parameter set ‘PB’ mimicking Priebe and Beuckelmann kinetics [21], and with the Fenton-Karma (FK) model [5] with modified Luo-Rudy I (MLR-I) parameters. In addition, we investigated how the methods perform when operating on a circular spiral wave core by applying them to the Aliev-Panfilov (AP) model [8].

All simulations were executed on a 2D isotropic square domain of myocardial tissue using Neumann boundary conditions and a 5-point stencil for the Laplacian. Integrating in time is done using forward Euler stepping, with values per model given in Table 1. There, we also present the typical APD values and other parameters that we use for the calculation of the phases.

**Table 1.** Overview of the performed experiments with relevant parameters.

|  | AP model [8] | BOCF model [9] | FK model [5] | hiAM monolayer (OM) [10] |
| --- | --- | --- | --- | --- |
| type | *in silico* | *in silico* | *in silico* | *in vitro* optical voltage mapping |
| grid size | $400 \times 400$ | $1200 \times 1200$ | $450 \times 450$ | $100 \times 100$ |
| $\Delta x$ | 1 | 0.25 mm | 0.262 mm | 0.25 mm |
| $\Delta t$ in ms | 0.01 | 0.009 ms | 0.163 ms | 6 ms |
| parameter set | as in [8] | PB | MLR-I |  |
| duration | 437.9 | 1250.01 ms | 815.31 ms | 12 282 ms |
| APD | 10.0 | 400.0 ms | 128.0 ms | 100 ms |
| parameters for LAT and $\phi^{\text{LAT}}$ | | | | |
| $V$ | $u$ | $u$ | $u$ | $u$ |
| $V_*$ | 0.5 | 0.3 | 0.65 | 0.75 |
| parameters for $\phi^{\text{act}}$ and $\phi^{\text{skew}}$ | | | | |
| $V$ | $u$ | $w$ | $u$ | $u$ |
| $R$ | $v$ | $s$ | $1 - v$ | $u$ delayed by 19.8 ms |
| $V_*$ | 0.5 | 0.6 | 0.65 | 0.75 |
| $R_*$ | 1.0 | 0.3 | 0.8 | 0.5 |
| $\phi_0/(2\pi)$ | 0.15 | 0.6 | 0.35 | 0.1 |
| $\phi_1/(2\pi)$ | 0.7 | 1.0 | 0.55 | 0.75 |

The spiral wave was generated by applying a classical S1S2 protocol: The western side of the square domain is initially excited to generate a plane wave. When the central point of the medium has finished repolarization, the south-western quarter of the domain behind the traveling wave is stimulated, such that a spiral wave (cardiac rotor) is formed. This protocol may also be applied rotated or mirrored.

Three frames of the first model variable *u* for each of the three chosen models resulting from numerical simulation were displayed in Fig 1.

#### 3.1.2 Optical voltage mapping experiments

To test our methods on *in vitro* measurements, in contrast to the *in silico* simulations, we used optical voltage mapping data derived from 10 cm^2^ monolayers of conditionally immortalized human atrial myocytes (hiAMs) following cardiomyogenic differentiation of these cells [10]. A voltage-sensitive dye is added to the culture, after which a real-time recording can be made of the intensity of emitted light, which is a measure of the local transmembrane potential. For the used recordings, pixel size was 0.25 mm and the sampling time between frames was 6 ms. The full protocols for cellular differentiation of hiAMs and the optical voltage mapping experiments can be found in the previous publication [10].

Gaussian smoothing with a kernel size of three grid points has been applied to each frame. The data have been rescaled such that each grid point has unit variance in time. Then, arbitrary units have been defined such that the resting state corresponds to optical activity *u* equal to zero, and the excited state to *u* = 1. To calculate the LAT phase *ϕ*^LAT^, we use a value of APD = 100 ms. This corresponds to a value in between the measured values of APD_50_ = (36.4 *±* 7.7) ms and APD_80_ = (136 *±* 12) ms [10]. Three frames of one of these recordings can be seen in Fig 1M-O.

As a second variable for calculating *ϕ*^act^ for this data set, we use the time-delayed version of *u* by 19.8 ms, which corresponds to roughly 15% of the duration of a rotation. The other parameters needed to calculate the phases can be found in Table 1.

### 3.2 A third phase definition

The aforementioned phase definitions *ϕ*^act^ and *ϕ*^LAT^ have each their downsides: Gradients in *ϕ*^act^ not only show PD but also wave fronts, and *ϕ*^LAT^ requires intense sampling over time of the medium. We propose to combine the advantages of *ϕ*^act^ and *ϕ*^LAT^ in a new phase, *ϕ*^skew^. This phase is designed as a computationally cheap approximation of the elapsed time phase *ϕ*^LAT^ that can be calculated using *ϕ*^act^. By construction, it does not require the whole history of *V*, but can instead be calculated using (*V, R*) at one point in time. In essence, *ϕ*^skew^ is a re-parameterization of *ϕ*^act^:

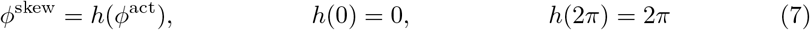

where *h* is furthermore monotonically rising. Essentially, the cycle visited by cells during the action potential is now labeled in a more free manner than the classical polar coordinates in the (*V, R*)-plane. As can be seen in Fig 4 by plotting *ϕ*^act^ vs. *ϕ*^LAT^, *ϕ*^LAT^ is also a re-parameterization of *ϕ*^skew^.

**Fig 4.**
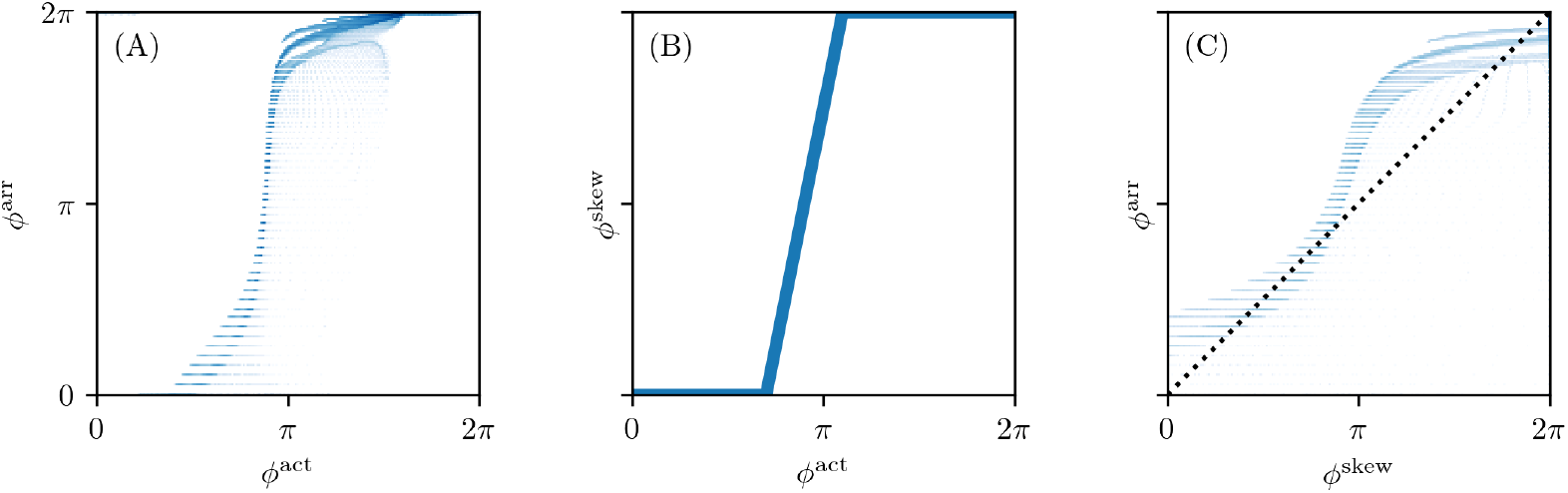
Correlation of the different phases. Phase data have been taken from the one snapshot in time of the Fenton-Karma simulation in Fig 3. Points for two phase values with darker shading correspond to higher logarithmic probability density. (A) The LAT phase *ϕ*^LAT^ stays close to zero while the state space phase *ϕ*^act^ is already increasing. It then transitions to 2*π* much quicker and stays relatively close to that value afterwards. (B) The skewed phase *ϕ*^skew^ is designed as a re-parameterization of the state space phase *ϕ*^act^ such that it approximates this behavior using a piecewise linear function. (C) The skewed phase *ϕ*^skew^ correlates more with the LAT phase *ϕ*^LAT^, getting closer to the dotted line where *ϕ*^skew^ = *ϕ*^LAT^.

In principle, one could fit a function *h* to a plot of *ϕ*^LAT^ vs. *ϕ*^act^, but we opt for a different approach here and define a piece-wise linear function *h*:

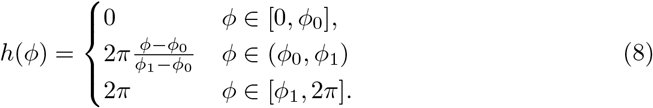

Values for *ϕ*_0_ and *ϕ*_1_ were manually chosen for the different reaction kinetics models used, see section 2.1 and Table 1.

Below, we will apply different PD detection methods on the three phases *ϕ*^act^, *ϕ*^LAT^ and *ϕ*^skew^, to see which performs best in visualizing PD structures.

### 3.3 Phase defect detection algorithms

#### 3.3.1 Requirements for phase defect detection algorithms

The aim of this paper is to provide and evaluate numerical methods that can be used as a successor of classical PS detection algorithms, but directed towards the detection of PDs instead. The following factors are taken into account when proposing the methods.

##### Zero or finite thickness

Due to the formation of a physical boundary layer (either by electrotonic effects or numerical smoothing, see [18]), a PD has a finite width in practice. Therefore, we see two options. A first option is to see the PD as an idealized structure with zero thickness, situated near the steepest spatial variation of phase or a spatial discontinuity in the LAT. A second option is to accommodate for the finite transition width, and describe the PD in a probabilistic manner, e.g. by regarding the phase gradient as a kind of PD density, below denoted as *ρ*. If desired, the PDL extent can then be determined by putting a threshold on *ρ*, a process which becomes easier if this density is normalized between 0 and 1.

##### Vertex-based or edge-based detection

Our algorithms take phase data on a set of nodes as input. We discriminate methods based on whether their output is on the nodes, edges or faces of the grid. Since the PDs have co-dimension one, it is natural to consider them as being situated on edges of the computational grid, either in 2D or 3D. However, the result of an edge-based method is not located on the original grid, such that methods that return values on the vertices of the grid (i.e. collocated with local phase data) can also be useful. Both edge-based and vertex-based methods are in contrast with PS detection: since PS have co-dimension 2, they are naturally calculated on the faces of the grid [5, 22]. Below, we provide for most PD detection methods an edge-based and vertex-based variant. We currently test our methods on a 2D Cartesian grid only and leave the extension to 3D and irregular meshes to future work.

##### Taking phase differences

Since phase is a cyclic variable, phase differences should be taken with care. Spatial derivatives are implemented in such a manner that an integer multiple of 2*π* is added in order to bring the result as close as possible to zero. Also trigonometric functions are adjusted such that they are indifferent to 2*π* differences. In some methods, the complex number *z* = *e*^i*ϕ*^ is calculated and the absolute value |*z*| is taken afterwards to obtain the phase instead of just using *ϕ*. This will make sure that a large jump in phase is not just attributed to the phase being cyclic.

##### Performance

In the results section (section 4), the different algorithms are compared in computational speed and relative performance. The location of PDs depends on the choice of algorithms for phase and PD calculation. To still make a comparison between methods possible, we designed an *in silico* experiment where the ground truth location of a PDL is known.

In the remainder of this section, nine different PD localization methods will be briefly presented. For the vertex-based algorithms, the output is a discretized scalar field: a non-negative PD density 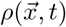 at time *t* that is defined in the points where phase is available to calculate the defect from (here either *ϕ*^act^, *ϕ*^LAT^ or *ϕ*^skew^). For the edge-based algorithms, the output is a number *σ*_*ab*_ computed from the pair of phases 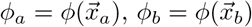 found at the vertices connected by that edge, which could be regarded as making up a vector field.

#### 3.3.2 Interpolation between vertex-based and edge-based methods

In what follows, 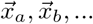 are positions of vertices *a* and *b*, 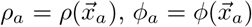 and *σ*_*ab*_ is a quantity defined on the edge of the mesh between 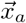 and 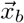. The set of neighbor vertices connected to vertex *a* is *𝒩* (*a*), containing *N*_*a*_ elements. In a 2D Cartesian grid, *N*_*a*_ = 4 inside the medium, but on the boundary of the domain or near obstacles, this value will be lower. In this case, the following formulas can still be applied with lower *N*_*a*_. In a 3D Cartesian grid, *N*_*a*_ = 6. See Fig 5 for graphical depiction of *ρ*_*a*_ and *σ*_*ab*_ on a portion of a Cartesian grid.

**Fig 5.**
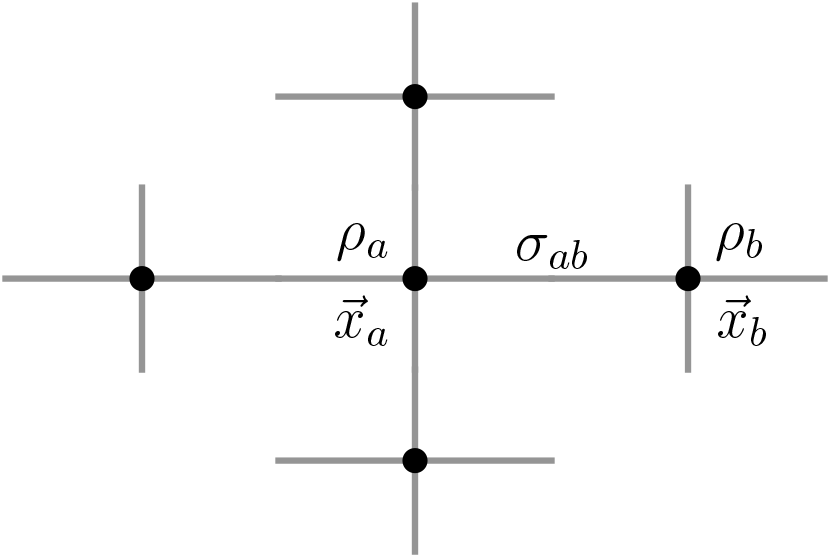
Overview of vertex- and edge-centered quantities. For vertices *a* and *b* at positions 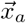 and 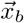, for which the phase *ϕ* has been calculated, we denote by *ρ*_*a*_ the vertex-centered PD density, and *ρ*_*b*_, respectively. For the edge *a* ↔ *b* between the two vertices *a* and *b*, we denote the edge-centered PD density by *σ*_*ab*_. The vertices that *a* is connected to are *a*’s neighbourhood *𝒩* (*a*).

If a quantity arises naturally along an edge (e.g. a gradient), it can be interpolated onto the vertex grid using

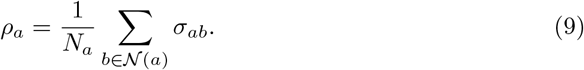

Conversely, if a quantity is found at vertices, it can be allocated to the edges using linear interpolation:

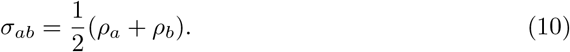

To distinguish between methods, the name of the method will be added in superscript, e.g. *σ*^CM^, *ρ*^PC^, etc.

#### 3.3.3 Overview of phase defect detection algorithms

In the following, we will present several different methods to detect PDs. A tabular overview of all the methods is given in Table 2.

**Table 2.** Overview of existing and proposed algorithms for PD detection.

| abbr. | name | rationale | centered | calc. time | source |
| --- | --- | --- | --- | --- | --- |
| CM | Cosine Method | Jumps in phase mod $2\pi$ are okay, in between: PDL. | on edges | 3.74 s | [15] |
| GLAT | Gradient of Local Activation Time | LAT jumps at PDL and wave front. | on edges | 2.13 s |  |
| RPG | Real Phase Gradient | Phase jumps at PDL. | on edges | 4.90 s | [16] |
| CPG | Complex Phase Gradient | Phase jumps at PDL. | on edges | 13.31 s |  |
| PC | Phase Coherence | Phase is not coherent at PDL. | on edges | 10.03 s |  |
| DM | Dipole Moment | Points on opposite side of PDL have opposite “charge”. | on vertices | 13.74 s |  |
| SVF | Spatial Vector Field | Rotation of gradient is close to zero, except at PDL. | on vertices | 8.37 s |  |
| AM | Angular Momentum | Use the classical topological charge also known as angular momentum. | on vertices | 1.47 s | [23] |
| IPM | Inflection Point Method | Change of sign of the phase Hessian. | on edges | 10.62 s |  |

Calculation times are for 500 frames in a medium of $450 \times 450$ pixels on an Intel Core i7-10875H processor in a Numpy implementation without specific optimization for speed.

##### Cosine method (CM)

Tomii *et al*. [15] introduced the following quantity to visualize PDs along an edge:

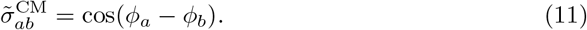

This method returns a value in [−1, 1], where low values indicate the presence of a PD. To derive a normalized PD density with values in [0, 1], we modify this to:

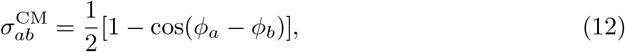

We define a vertex-based version of this quantity via Eq (9), which we denote with 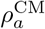.

##### Gradient of local activation time (GLAT)

It is expected that around a PD, the LAT does not vary smoothly but instead jumps across this line. This implies that the gradient in the neighbourhood of the PDL should be much larger than further away where neighbouring vertices are activated subsequently.

This relation is easily expressed using edges:

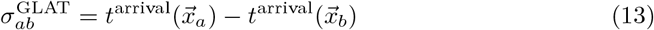

Note that we are using a second condition here: If the elapsed time since excitation of either vertices is exactly zero, we still set 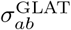 to zero. The rationale here is that the jump is due to the wave front passing instead of pointing to a PD.

A vertex-based variant is found by averaging over all edges leaving the same vertex, see Eq (9), applied to the absolute value of the LAT difference:

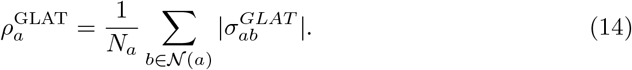

##### Real phase gradient (RPG)

In previous work [16], we considered phase gradients, disregarding 2*π* phase differences:

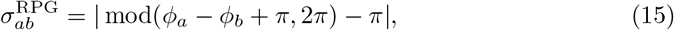

returning values in [− *π, π*]. The normalized cosine method (Eq (12)) can be seen as a mapping of this interval to a PD density taking values in [0, 1].

##### Complex phase gradient (CPG)

The next method works in a similar fashion, but to avoid the modulo operation, we look for gradients in the complex number *z* = e^i*ϕ*^:

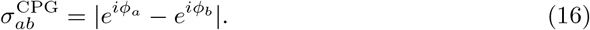

From this a vertex-based density can be computed using Eq (9).

##### Phase coherence (PC)

Inspired by the literature of phase oscillators [24], we define the phase coherence of a vertex *a* with neighbours *b* as:

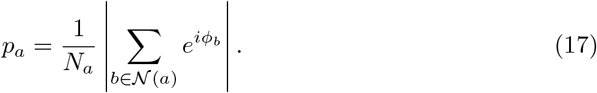

The PD density is then defined as

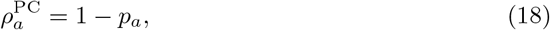

such that *ρ* is large when the coherence is low, since a low phase coherence is expected near a PDL. This index *ρ*^PC^ is normalized in [0, 1] with large values indicating high PD probability.

Note that applying the PC method to only two vertices connected via an edge delivers

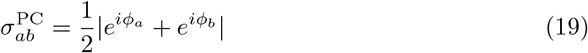

returning 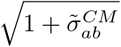.

##### Dipole moment (DM)

Along a PD, the phase values that surround a given point are expected to be divided into two groups, one on either side of the PDL. We could perhaps detect this splitting by using the concept of the dipole moment of a charge distribution, where the complex number *z* = e^*iϕ*^ takes the role of charge:

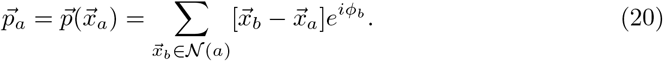

The point-based PD density is then found by taking the norm of this complex vector:

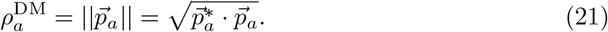

where *\** denotes complex conjugation.

When two vertices are connected by one edge, the edge-based implementation will recreate the CPG method. For this reason, no direct implementation of the latter was done. Still, edge-based values can be computed via interpolation, see Eq (10).

##### Spatial vector field (SVF)

When Stokes’ law is applied to the expression of topological charge, one finds

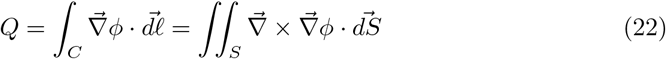

where *C* is the boundary curve to the region *S*. Since for a continuous field *ϕ*, the rotation of the gradient 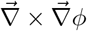 vanishes everywhere, a continuous phase field cannot bear non-zero topological charge. Nevertheless, computing *Q* for all faces of the grid has been used to find PSs [3].

Inspired by the right-hand side of Eq (22), and replacing *ϕ* by *z* = *e*^*iϕ*^ to get easier differentiation, we propose:

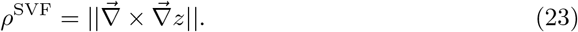

The motivation for this method is that a PD is essentially a discontinuity in the field *ϕ*. At such a discontinuity the rotation of the gradient may be different from zero. In our current implementation, we calculate the gradient in a vertex-based manner, e.g. ∂_*x*_*u*(*x, y*) = [*u*(*x* + *dx, y*) − *u*(*x* − *dx, y*)]*/*(2*dx*), such that the result is also vertex-based.

##### Angular momentum (AM)

A classical method to detect the central region in a spiral wave is using the pseudo-vector: [23]

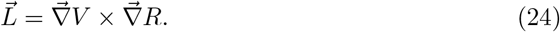

Far away from the spiral core, the activation resembles a plane wave, making 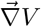 nearly parallel to 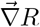, such that 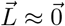 except near the core of the spiral. Since PS can be seen as a limit of a PDL with vanishing length, we will visualize

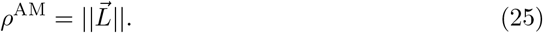

An edge-based method can be derived by taking:

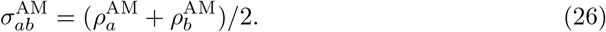

##### Inflection point method (IPM)

Given that a PDL in practice connects two regions of different phase in an abrupt but continuous manner, it is interesting to look where the phase transition is the steepest, and localize the PD there.

For 1D functions, an inflection point is found where *f* ^*′*^ (*x*) changes sign. This can be translated to the condition *f* ^*″*^ (*x*) = 0. To find the same region for a 2D function, we express that we want an inflection point when stepping in the direction of the local phase gradient. With 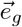 as the normalized gradient vector:

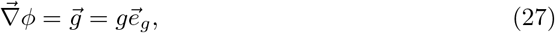

the spatial derivative in the gradient direction is 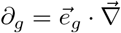. With this, ∂_*g*_*ϕ* = *g*, and the concavity in the direction of the gradient becomes:

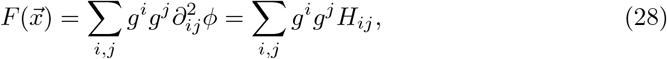

where the Hessian of the phase is 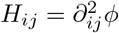. Hence, the PD can be found as the set of points where 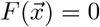. This method is unlike the mentioned algorithms above, since it immediately returns a line, i.e. PDL of zero thickness. Note that in practice, one needs to impose a minimal value of ||*ρ*^*GLAT*^|| such that the background region with low PDL density is filtered out.

To compare this method to the other algorithms, we color the edges where 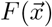 changes sign with the phase gradient along that edge, i.e.

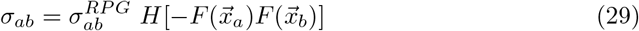

with the Heaviside function *H*, which takes the value 1 for inputs larger than zero and 0 otherwise.

### 3.4 Visual representation of the methods

For the methods that return a vertex-based PD density *ρ*, we simply color the pixels in the rectangular grid according to *ρ*. For the methods that return an edge-based PD indicator *σ*, we color the dual grid, i.e. we color every point in the plane according to its nearest edge. This results in a coloring of the plane using pixels that are 45^*°*^ tilted and centered around the midpoint of edges in the original grid. In this way, interpolation between edges and vertices does not affect the presented results.

### 3.5 Post-processing of phase defects

Having obtained a PD density 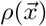 using one of the methods, we keep only points above a threshold value *ρ*_*c*_ to obtain a set of points on the PDL, and connect it using the minimal spanning tree graph algorithm. Thereafter, the smallest branches of each tree are cut to gain a discrete representation of a PDL, centered at the vertices of the image grid.

To measure PDL length *L*, the PDL points are connected by line segments; the sum of their lengths is taken as an estimate to the PDL length.

To measure PDL precession speed, we first selected a spatial region where only one PDL was seen during the timespan of interest. Then, we performed principal component analysis (PCA) to the point cloud of the PDL at all time instances to obtain the main vector of alignment 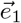. The angle between this vector and the positive *x*-axis is taken to be *β*, after which linear regression of *β*(*t*) = *β*(0) + *ωt* yields an estimate for the precession frequency *ω* and the precession period 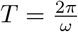.

### 3.6 Performance tests

#### 3.6.1 Data sets with Gaussian additive noise

The data sets we are working with in this paper are quite smooth in the sense that there is very little noise in them. This is due to the data from *in silico* simulations being smooth by design and the optical voltage mapping data being pre-processed with Gaussian smoothing before calculating the phase and phase defects (section 3.1.2).

To gauge how well the algorithms are able to deal with noise, we have added noise following a normal distribution with different signal to noise ratios (SNRs) to the data sets.

In the context of our work, we define SNR as the ratio of the standard deviation of the signal *u* to the standard deviation of the noise *n*

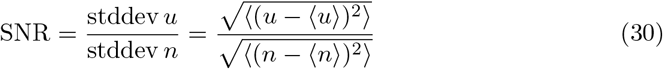

with 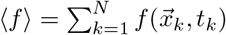 on all *N* points of the space-time grid.

#### 3.6.2 Data sets at lower spatial resolution

In many contexts, the only available data are at much lower spatial resolution than the *in silico* simulations and optical mapping experiments considered in this work. For instance, there are electrode arrays that are inserted via balloon catheters with 64 electrodes [25].

To still be able to assess the effectiveness of the phase and PD algorithms at such resolutions, we down-sample our data by a factor of *N* ∈ N by pooling *N × ℕ* grid points together using the arithmetic mean. Vertices that are not inside the medium are not included in this mean.

#### 3.6.3 Data sets: recovery of an obstacle

There is no ground truth in the exact location of the PD as it depends on the choice of algorithms for phase and PD calculation. Still, experiments are possible where the expected location of a PDL is known. We have conducted such an experiment *in silico* where we add an elongated, thin obstacle for a rotor to attach to.

For this, we have taken the last frame from the experiment using the FK model as the initial state of this experiment and placed such an obstacle at the core of the observed rotor.

For the length of the obstacle, we have chosen roughly the previously observed PDL length. We choose its width to be less than eight grid lengths 8Δ*x*, such that when we downsample the data by a factor of eight as in section 3.6.2, all pixels in the medium are again active. To put it briefly, the obstacle is designed such that it is too thin to be detected directly at that lower spatial resolution. Instead, we use phase defect detection to recover the obstacle.

For this experiment, we used the parameters for the FK model as in Table 1, except for the time step Δ*t* = 0.1 ms and duration 800.1 ms.

To recover the obstacle from the simulation, we have used the different phase and PD algorithms to calculate PD densities 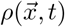. Recall that all methods are designed such that high 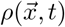 corresponds to high likelihood of a PD being located there. We calculate a prediction of the location of the obstacle 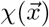 based on 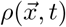 as follows:

1. For a duration of at least one rotation of the spiral, calculate the mean value of the PD density 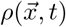 at each point in space. Call this quantity 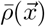.
2. As 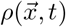 is close to zero except for at the phase defect, the distribution of values in 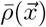 will also be heavily skewed towards lower values. Therefore, we clip 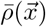 to the interval from the 40th percentile to the 99th percentile to get rid of both small fluctuations around zero and outliers.
3. To obtain 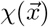, we finally rescale to the interval [0, 1] and then round to *{*0, 1*}*. This quantity can be thought of as an approximation of the characteristic function 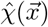 of the set of points in the obstacle.

Finally, to be able to judge how well the obstacle has been recovered, we calculate the classification error as the fraction of misclassified points by the prediction 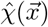 with respect to the ground truth 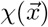.

## 4 Results

We here apply the different proposed detection methods for the three phases *ϕ*^act^, *ϕ*^LAT^, and *ϕ*^skew^. We do this in three cardiac monodomain models and compare performance of the methods (section 3.1.1). Finally, we apply a selection of methods to an experimental dataset obtained by optical voltage mapping of a monolayer culture of cardiac cells (section 3.1.2).

### 4.1 Comparison of different phase definitions

Fig 3 shows the three phase definitions applied to a snapshot of the three monodomain models, the Aliev-Panfilov (AP) model [8], the Fenton-Karma (FK) model [5], the Bueno-Orovio-Cherry-Fenton (BOCF) model [9], and an optical voltage mapping experiment.

The AP model shown in the first row of Fig 3 produces a rigidly rotating spiral. With *ϕ*^act^ and *ϕ*^skew^, a PS is seen. However, due to the thresholding on *V* used to determine LAT, the inner part of the core region is never excited, such that *ϕ*^LAT^ shows an abrupt change at the trajectory of the classical PS, which will be picked up as a PD below.

In the simulations with linear core (FK and BOCF models), *ϕ*^act^ shows sudden transitions at the rotor core and the wave front, while *ϕ*^skew^ and *ϕ*^LAT^ only show a distinct phase gradient near the conduction block line.

The optical voltage mapping experiment in Fig 3J-L shows apparent PSs for *ϕ*^act^ and *ϕ*^skew^, but an extended PD for *ϕ*^LAT^. Hence, at first sight, it resembles the AP spiral, but this relation will be further investigated below using the PD detection techniques outlined above.

Fig 4 shows a scatter plot between the different phases for the FK frame shown in Fig 3. We took the convention that the phase at the resting state is 0. The skewed phase *ϕ*^skew^ with parameters tuned as outlined above resembles the elapsed time phase *ϕ*^LAT^ much closer than the state space phase *ϕ*^act^. In short, *ϕ*^skew^ is an approximation to *ϕ*^LAT^ that does not require observation of the system during the previous excitation sequence.

### 4.2 Comparison of phase defect detection methods in simulations

We have applied all PD detection methods (section 3.3) to all experiments, for all phases. For definiteness, we only show the result for the FK model in Fig 6 and 7, but the others can be found in the Supplementary Materials (S1 Fig, S2 Fig, S3 Fig, and S4 Fig).

**Fig 6.**
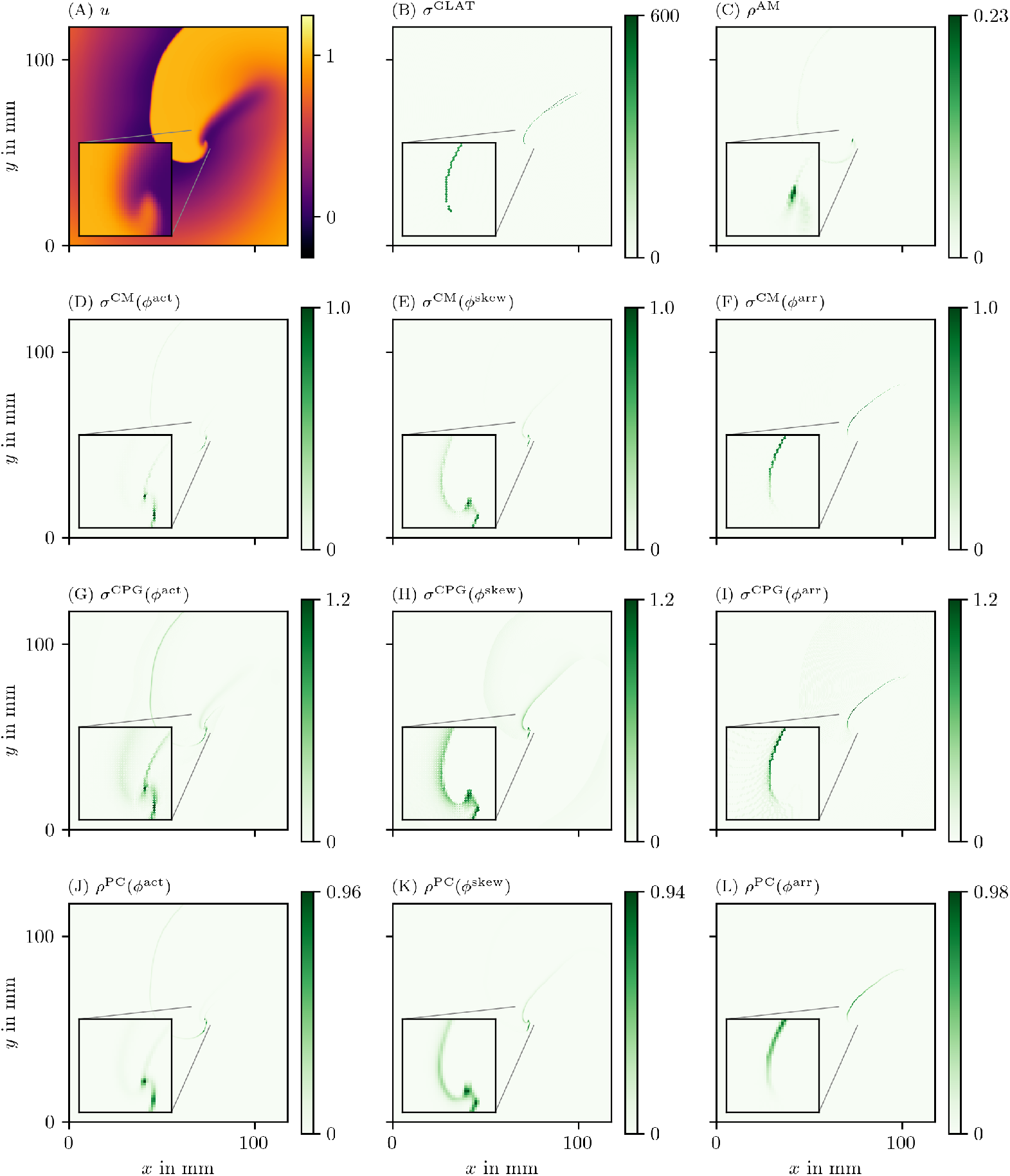
Overview of PD detection methods for one snapshot of the FK data set. The PD on vertices *ρ* or edges *σ* is measured in arbitrary units. The same coloring as in Fig 2C is used here. As the PDL has a width of only a few grid points, we zoom in around the turning point to get a better view of the structure on the grid. For reference, we also show the corresponding frame of the transmembrane voltage *V* = *u* in panel A.

**Fig 7.**
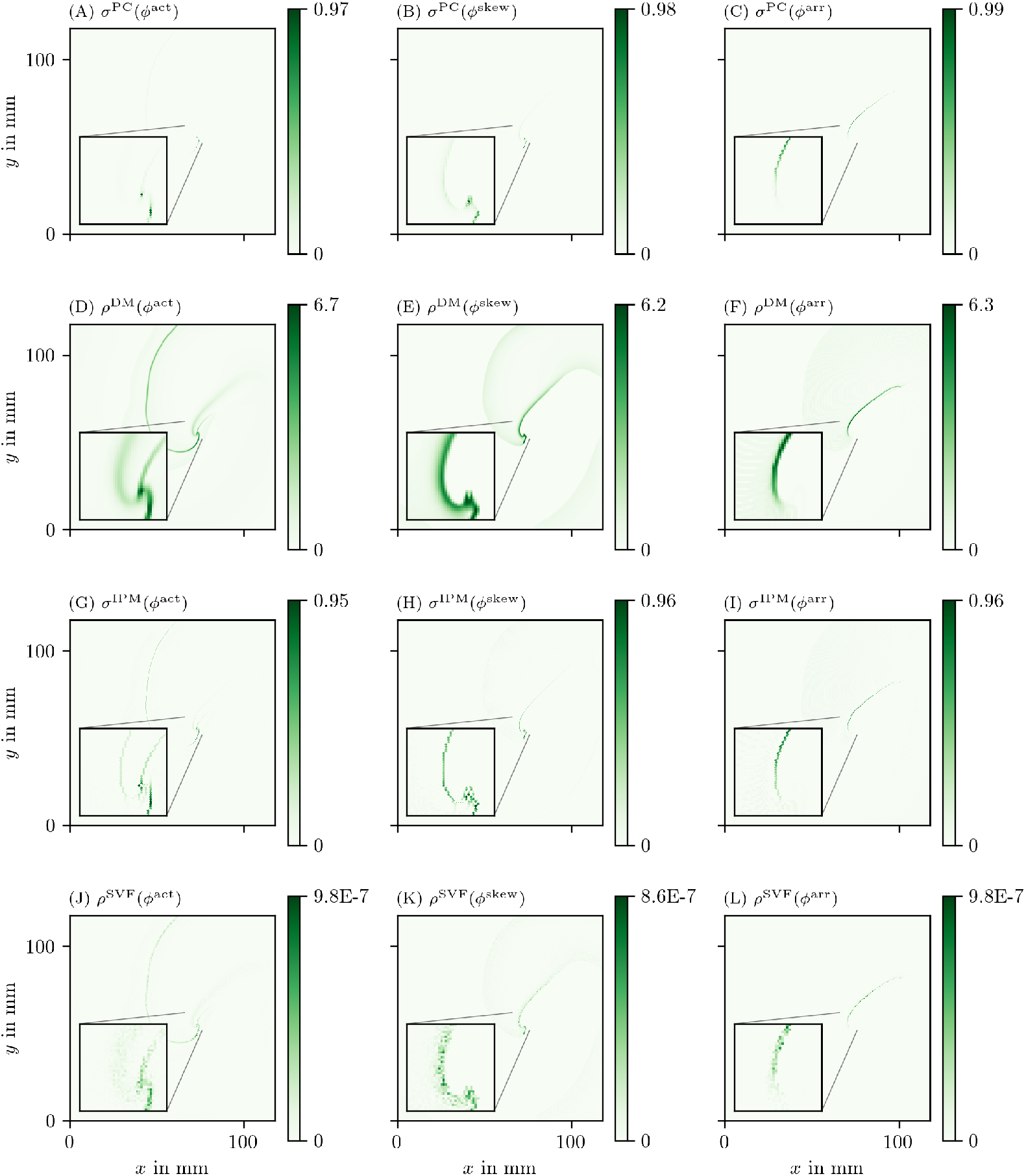
Overview of more PD detection methods for one snapshot of the FK data set as in Fig 6. Each row shows a detection algorithm, applied to the three different phase definitions.

In the following, we will present a selection of the results exhibiting the common features and our main observations regarding the different methods.

In general, all methods return low densities away from the wave front and PDL and higher values near the region of interest, although the precise PD density distribution is different between the methods.

In Fig 6B, the GLAT method clearly shows the conduction block line at the rotor core. Since LAT is discontinuous there, the set of points is thin such that small gaps can be seen. At the rightmost part, the PDL doubles, since the process of reaching and leaving the rightmost turning point, both leave a discontinuity in LAT. Moreover, the PDL’s precise location depends on the chosen threshold *V*_*x002A;*_.

The AM method (Fig 6C) locates only the site where the wave front meets on the PDL. Also it can be seen that *ρ*^AM^ is located at the wave front, though with much lower magnitude. This is consistent with this method traditionally being used for PS detection.

The other methods in Fig 6-7 are phase-based. In each case, the wave front is most clearly seen as an artifact using *ϕ*^act^, less visible using *ϕ*^skew^ and absent in *ϕ*^LAT^.

The CM, RPG and CPG methods (Fig 6D-L) and PC and DM methods (Fig 7A-F) give all qualitatively similar results: With *ϕ*^act^ and *ϕ*^LAT^, not only the PDL but also the end point of the wave front (tip) is stressed. The *ϕ*^LAT^-variant distinctly shows the PDL, as the wave front is filtered out by the definition of *ϕ*^LAT^.

The IPM method shows a line that is only one pixel wide, as it was designed to localize the PD at the site of steepest phase variation.

Finally, the SVF method yields many points in the region of interest, but the result is noisy even in this idealized simulation.

### 4.3 Comparison of phase defect detection methods in an optical voltage mapping experiment

We also applied the different phase definitions and detection methods to the excitation sequence observed in a hiAM monolayer, as detailed in section 3.1.2. In Fig 8 and Fig 9, we show the results of this process for all of those methods. Fig 8A shows the optical intensity at a given time in a multiple-spiral state. The non-phase methods GLAT and AM in Fig 8B-C show non-zero densities at several positions that are similar in both methods. The four most intense points, at which either a PS or PDL could be present, are confirmed by the other methods (CM, RPG, CPG, PC, DM, IPM) using *ϕ*^act^ and *ϕ*^skew^. When using *ϕ*^LAT^, the same 4 points are prominent, but they extend to a line (PDL) since the LAT and *ϕ*^LAT^ keep track of the recent history of excitation. Compared to the simulation data, more background structures are seen in the optical voltage mapping data, such as borders of excited regions and a staircase effect in LAT due to the time sampling.

**Fig 8.**
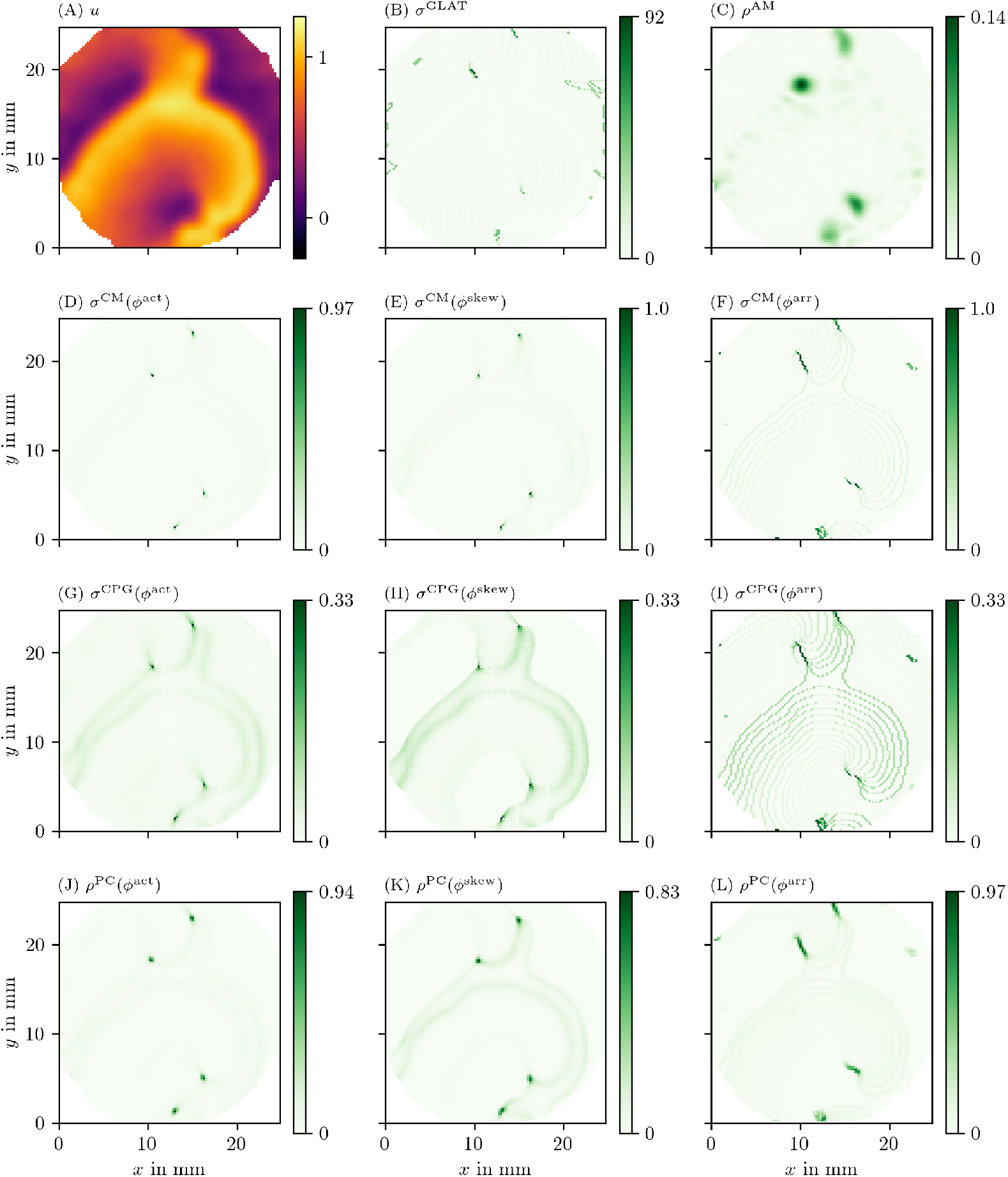
Overview of PD detection methods for one snapshot of the optical voltage mapping data. The data are presented in the same way as in Fig 6.

**Fig 9.**
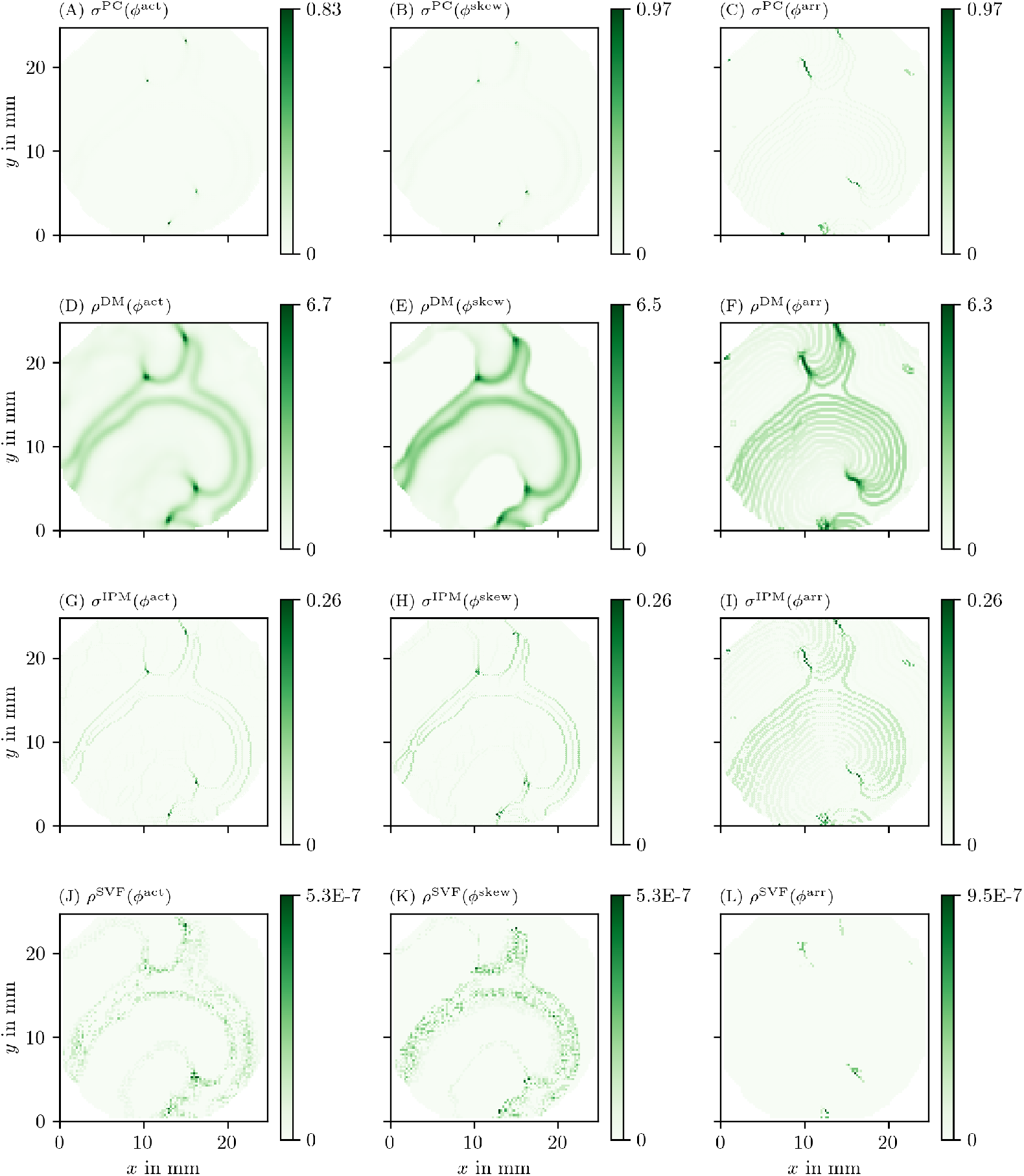
Overview of more PD detection methods for one snapshot of the optical voltage mapping data as in Fig 8.

### 4.4 Properties of PDLs *in silico* and *in vitro*

The presented methods allow to characterize the observed PDLs in terms of length *L* and orientation angle *β*, which is a further step in the quantitative analysis of excitation patterns.

Fig 10 shows the length over time of one PDL, in simulations and experiment. The PDL is detected using the LAT phase *ϕ*^LAT^ as input for the phase coherence method (PC). We observe that the PDL length varies over time. Its time-averaged value is summarized in Table 3.

**Fig 10.**
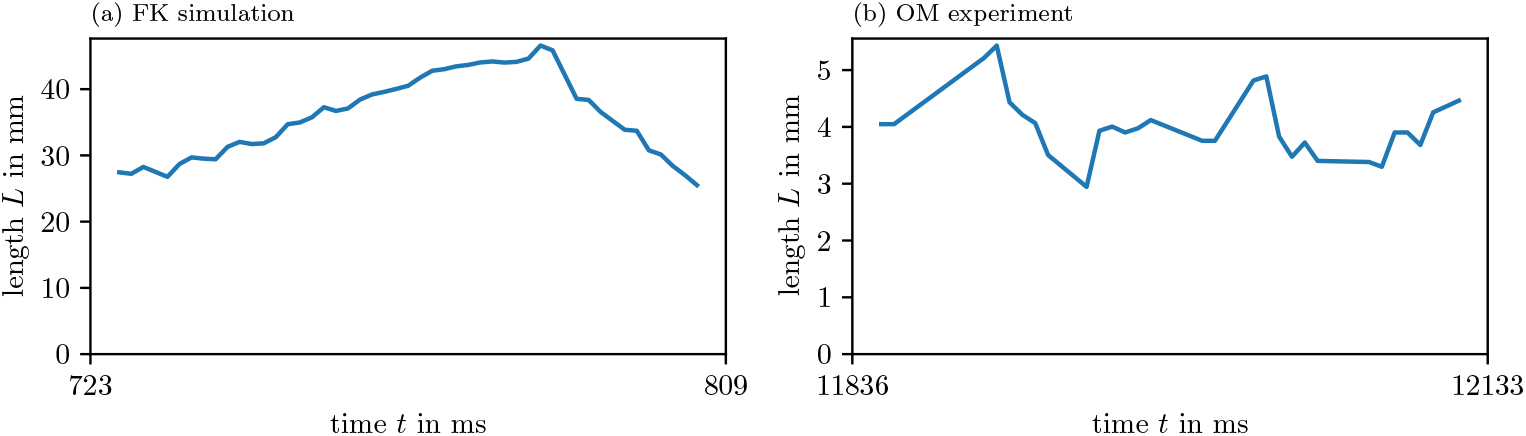
Length of detected PDLs over time. For one of the PDLs detected by the PC method for *ϕ*^LAT^, we show how its length changes over time. This length fluctuates around an average value.

**Table 3.** Statistics of one PDL’s length and precession over time.

| model / experiment | average length $L$ | precession period $T$ |
| --- | --- | --- |
| AP simulation | $(15.07 \pm 0.67)$ | $(33.123 \pm 0.011)$ |
| FK simulation | $(35.9 \pm 6.2)$ mm | $(1223 \pm 29)$ ms |
| BOCF simulation | $(76.1 \pm 7.8)$ mm | $(57\,508 \pm 87)$ ms |
| hiAM optical voltage mapping | $(4.01 \pm 0.55)$ mm | $(169.2 \pm 9.3)$ ms |

We observe similar values for other PDLs in the data.

In both simulations and experiment, we also estimated the precession period *T* of the PDL, see Table 3 and Fig 11. The orientation of the PDL changes almost linearly in all cases, hence, we observe quite low variance along the fit linear functions. On the one hand, in the FK and BOCF simulations, we see that *β* almost stays constant, but slightly precesses in one direction. On the other hand, in the AP model and optical voltage mapping data, the precession takes place in a much shorter period of time: *T* is just 1.5 to 3 times longer than the APD in both of those cases.

**Fig 11.**
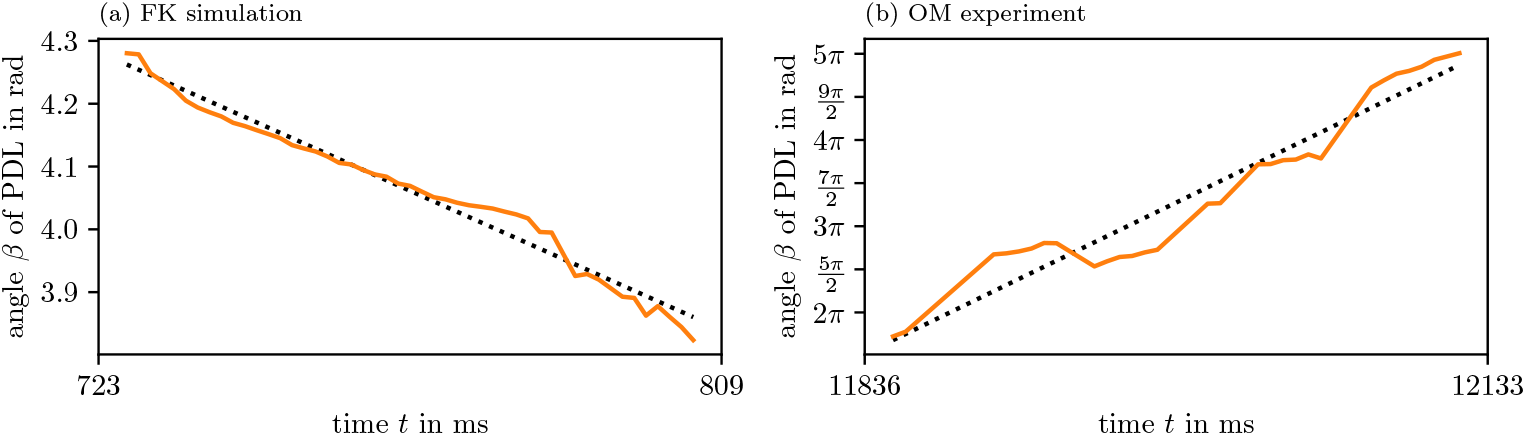
PDL orientation over time. For these figures, we use the same PDLs as in Fig 10 and Table 3.

**Fig 12.**
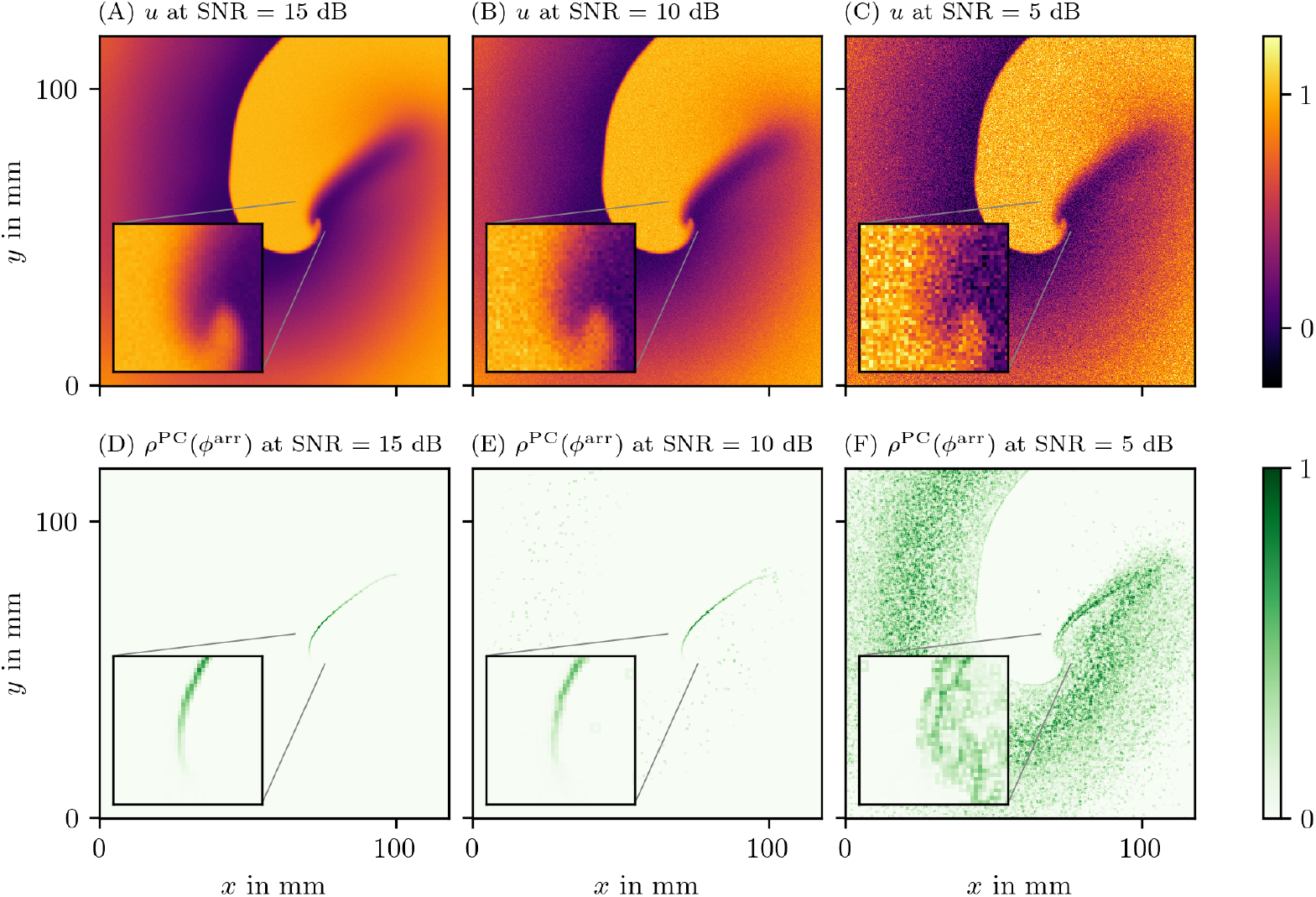
Effect of additive white Gaussian noise on the detection of PDs in the FK data set. The PD has been determined using the PC method applied to the LAT phase *ϕ*^LAT^ with two thresholds 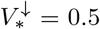 and *V* = 0.65. In the three columns, we vary the SNR. The data are presented in the same way and same point in time as in Fig 6.

### 4.5 Robustness to noisy data

We have also investigated how the methods perform under noisy conditions. For this we considered noisy data which were obtained as outlined in section 3.6.1. We then ran the different PD detection algorithms on those sets as well.

A general observation is that if the input data *V* is noisy, so are the state space phases *ϕ*^act^ and *ϕ*^skew^, which then can be seen in *ρ* as well.

Another effect can be observed for *ϕ*^LAT^, as it is based on LAT: When *V* increases such that it crosses the threshold *V*_*\**_, the value of LAT is updated to the current time. This can be triggered by noise which is especially critical right after *V* decreases falling below the threshold. A random fluctuation due to noise can then push it above the threshold again.

To counter this effect, we use a second threshold value 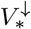 in the calculation of LAT and *ϕ*^LAT^ (section 2.1). A good value for 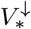 can be obtained by decreasing *V*_*\**_ by an offset that is proportional to the SNR. This value also depends on the wave back in the tissue model.

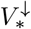 must be chosen low enough such that it suppresses LAT triggering due to noise at the wave back, but high enough that the tissue always repolarizes below it before the tissue can be excited again.

As an example, we show the PD *ρ* as determined by the PC method based on the LAT phase *ϕ*^LAT^ for the FK data set at three different levels of noise in Fig 12. We choose 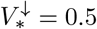 and *V*_*\**_ = 0.65. It can clearly be seen that this method succeeds to locate the PDL for SNRs above a critical value. In this example this critical value is around 10 dB. Here it can be seen that at various pixels in the medium a random fluctuation due to noise has pushed the input data from below 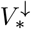 to above *V*_*\**_. At this stage it is still quite clear where the PDL is located. With more noise than this, however, this effect takes over. This leads to being unable to distinguish the PDL from the noise artifacts in the very noisy case with SNR = 5 dB.

### 4.6 Performance at lower spatial resolution

We also applied the PD detection algorithms to the data sets at different, lower spatial resolutions (section 3.6.2). In Fig 13, we present a frame of the input data *V* and the phase defect *ρ* for PC method and the LAT phase *ϕ*^LAT^. It can be seen that even at those resolutions, the PDL can still successfully be identified. For grid lengths larger than the length of a PDL, only few pixels have high enough *ρ* to be considered a PD. This illustrates that PSs and PDLs can not be distinguished from one another when resolution is too low.

**Fig 13.**
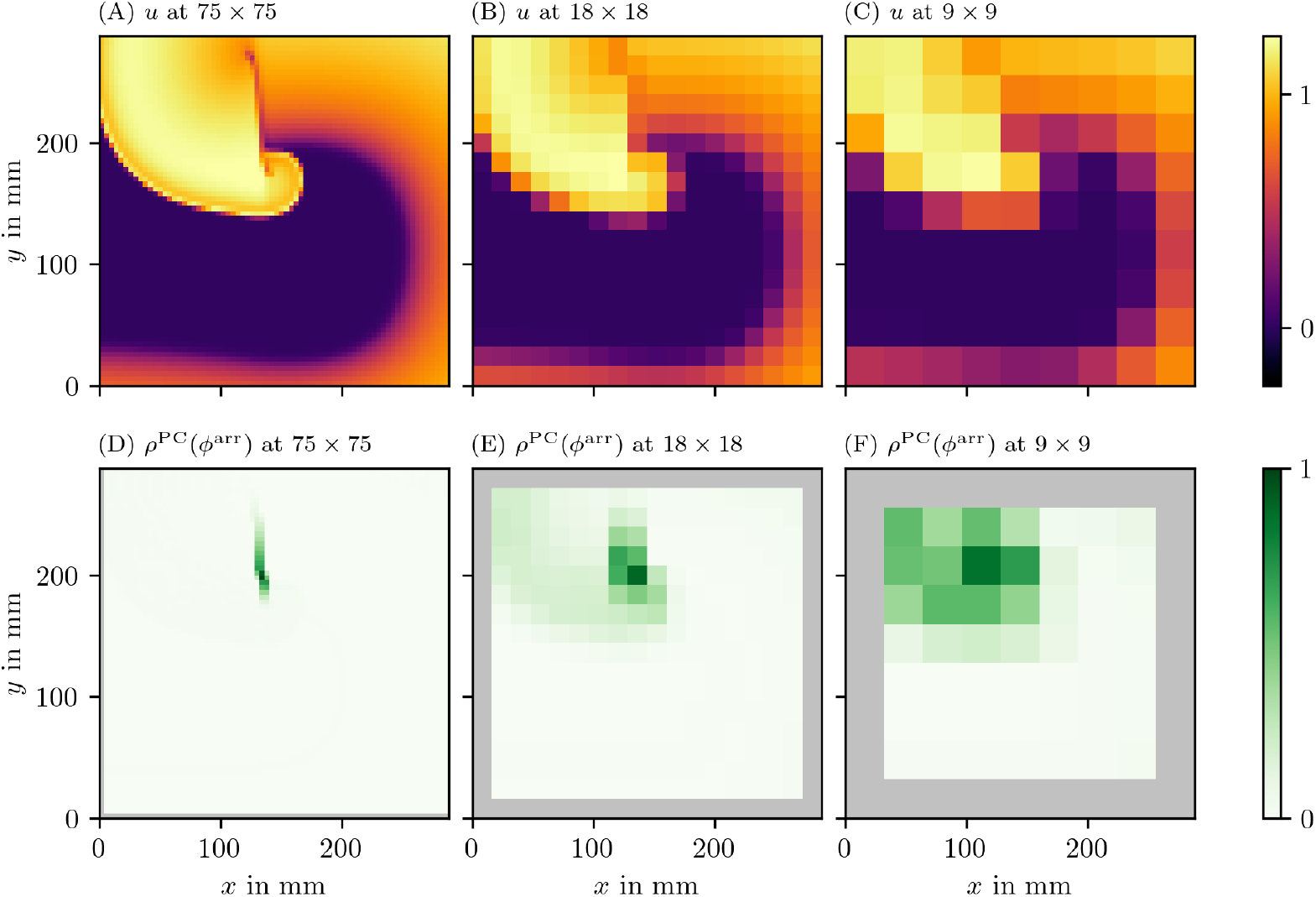
Performance of PD detection at different spatial resolutions for the BOCF data set. The PD has been determined using the PC method applied to the LAT phase *ϕ*^LAT^. The data are presented in the same way as in Fig 6.

Also note that the jump in phase due to the wave front and back passing through an area is larger at lower spatial resolution. As a PD is a large jump in phase, our methods detect this jump as well. The wave front is therefore harder to distinguish from a PD at low resolutions. This effect is much stronger when using *ϕ*^act^ but can be reduced by using *ϕ*^LAT^ or *ϕ*^skew^.

### 4.7 Recovery of an obstacle

In section 3.6.3, we designed an experiment such that a ground truth location for a PDL is known. An elongated obstacle was placed such that a rotor could attach to it.

Looking at the resulting data, we see that in fact the rotor has attached to this obstacle. A PDL formed around the obstacle.

With these recordings in *u* and *v* for the FK model, we moved on to calculate the phases *ϕ*, PD densities *ρ*, and the approximation of the characteristic function *χ* of the obstacle for all different methods. In Fig 14 and Fig 15, we present the recovered obstacles by each of the methods.

**Fig 14.**
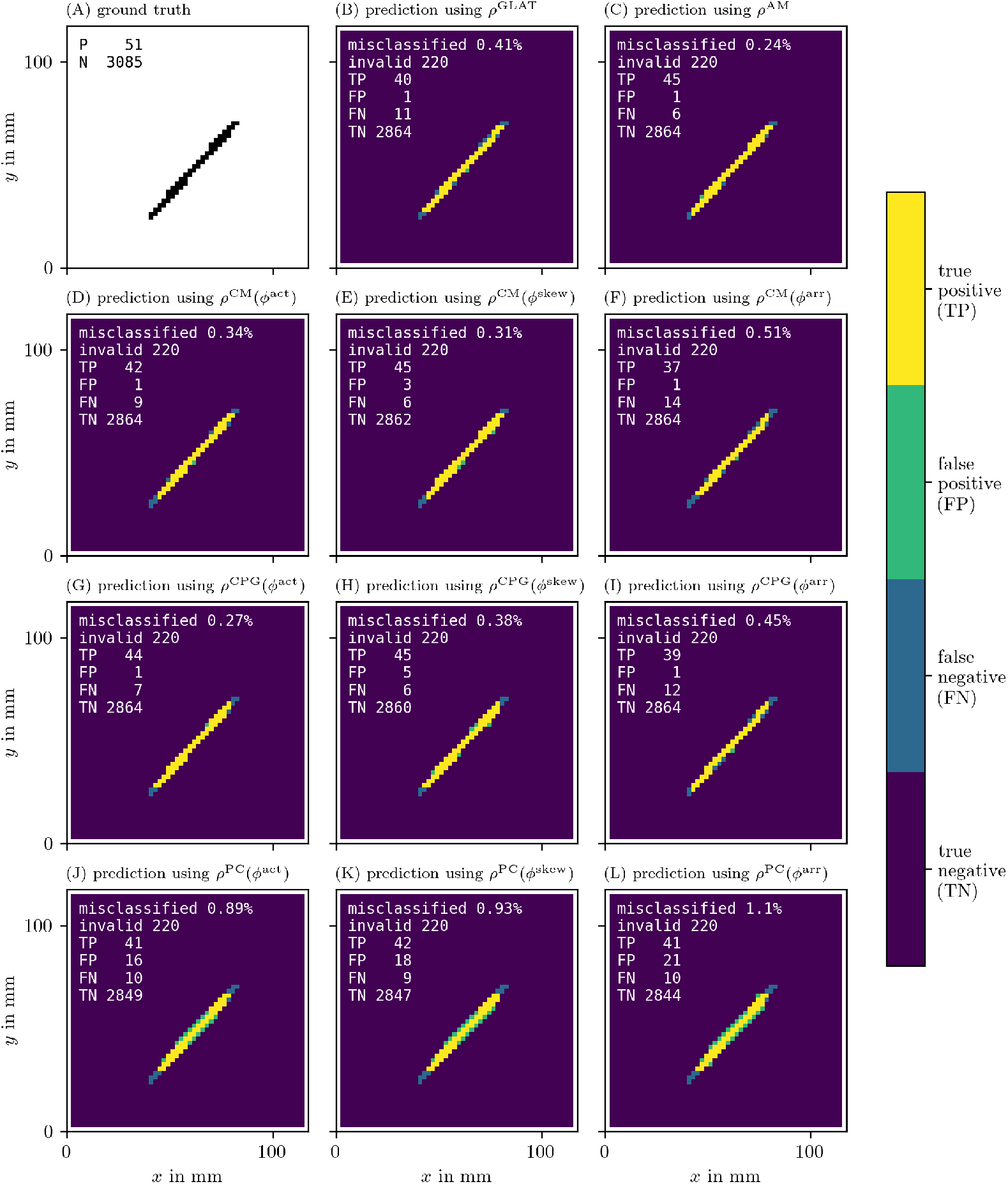
Recovery of the location of an obstacle based on each of the PD detection methods. As outlined in section 3.6.3, we calculate an approximation of the characteristic function 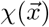 of the obstacle for each of the methods. We then calculate the classification error comparing this prediction to the ground truth 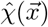.

**Fig 15.**
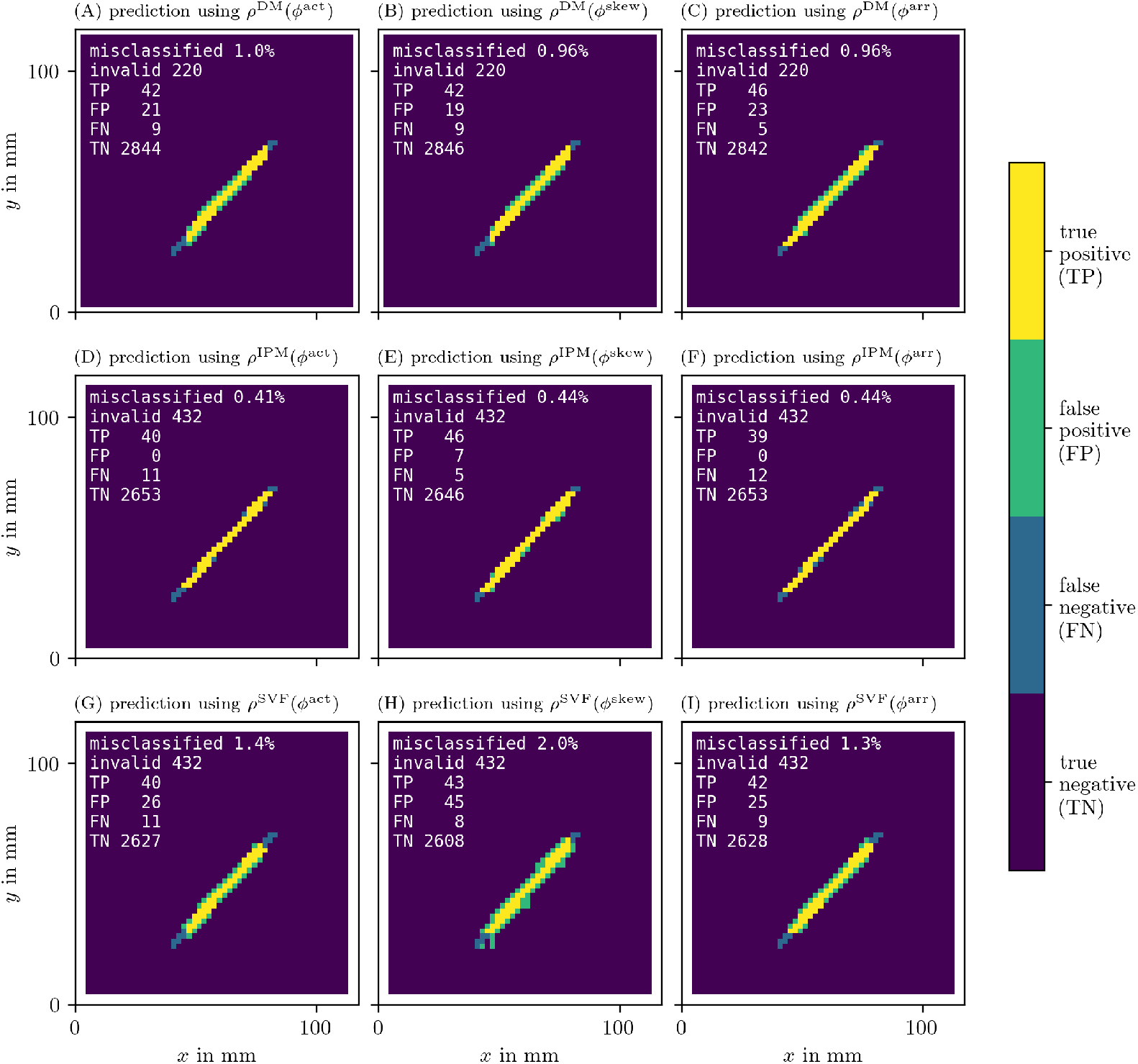
Recovery of the location of an obstacle based on each of the PD detection methods as in Fig 14.

It can be seen that all methods perform well at recovering the obstacle leading to a classification error of around or less than 1 %. We have therefore validated the methods and shown that they are able to locate PDLs.

The AM method is also able to predict the obstacle well, even though it is based on PSs. This is because PSs follow the PDL. While the PD based methods show the full extent of the line, the PS based AM method only returns a point-like region.

While there is only functional re-entry in the optical voltage mapping data set, we can still use the same algorithm to recover the long-term location of the PDL in the OM data using the PC method and LAT phase. As can be seen in Fig 16, the cores of the spirals can successfully be detected using this method. Our prediction 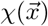 of the sites of functional obstacles via PDLs can be compared to so-called driver domains, specifically to driver-density maps which are based on PSs [26].

**Fig 16.**
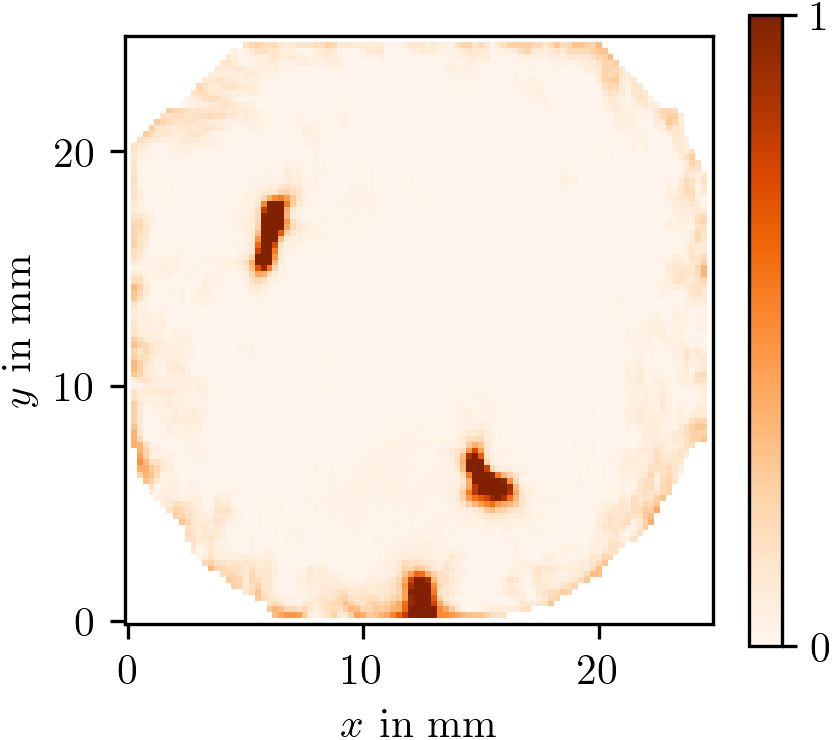
Prediction of the characteristic function *χ* of the effective obstacles based on *ρ*^PC^(*ϕ*^LAT^) for the OM data set. Due to the functional re-entry, the PDLs in the centres of the spirals observed in this recording effectively form functional obstacles. These differ from anatomical obstacles in the way that functional ones could excite and do in fact excite in this case before the rotors form. We recover those obstacles via *χ* as outlined in section 3.6.3.

## 5 Discussion

In this paper, we provide and compare several numerical methods to detect a PDL, a recently proposed structure present at the core of a rotor as an alternative for the classical PSs [15, 16]. Here, we attempt to improve the simple PDL detection methods from these works (CM & RPG) and tested them on simulations and experimental data.

Several phase-based algorithms were applied, not only the *classical phase ϕ*^act^ but also the recently introduced LAT-based *LAT phase ϕ*^LAT^ [16], since the LAT better keeps the spatio-temporal activation and therefore more clearly shows extended PDs. In addition to a systematic comparison between detection methods, we also introduce a third phase, the *skewed phase ϕ*^skew^ in this work. *ϕ*^skew^ was designed as a way to estimate *ϕ*^LAT^ from a single snapshot. This is useful in the post-processing of data from experiments or simulations, where a sparse time-sampling was used.

As *ϕ*^act^ transitions on small scales at the wave front and back, they may wrongly be identified as PDLs for this phase definition. Although *ϕ*^skew^ filters the wave front and back better than *ϕ*^act^, we still find that direct measurements of LAT and *ϕ*^LAT^ produce better-resolved PDLs (cf. Fig 6). Still, in the regime of fast depolarization, wave front and wave back can be considered PDs. These can be distinguished from PDLs due to conduction blocks via two criteria:

1. Wave fronts propagate in space.
2. Wave front and wave back connect always the same phase values, while PDLs connect other values.

When comparing the different methods to convert phase into a PD, we find a good performance and strong correlation between the previously coined cosine method [15] and the real phase gradient method [16]. Both methods are based on the same idea (measuring angular differences along a circle) and therefore the correlation comes as no surprise. Qualitatively similar performance is found by related methods (CPG, CM, PC). The IPM method also works well. Some advanced methods such as DM, and SVF actually performed worse in terms of contrast and noise suppression. Of special interest is the classical AM method [23], which consistently finds the end point of the wave front, even if lying on a PDL, and the GLAT method. The gradient of LAT correctly identifies the PDL and locates it very sharply (by construction); however, its precise location depends on the chosen threshold (*V*_*\**_) to classify tissue as excited or not. Regarding the calculation time, we find that the CM, AM and GLAT are the fastest and therefore recommended to use for processing larger datasets, e.g. extended in time or in three spatial dimensions.

To calculate the state space phase for the OM data, we have used a time-delayed version of the observed variable *u* to be able to calculate the state space phase *ϕ*^act^. Note that this traditional approach is unable to detect PDLs, as it gives distinct points where *ϕ*^act^ is high, corresponding to classical PSs. This explains why line shaped PDs were not investigated closer before. In panels Fig 8FIL and Fig 9CFIL that use the LAT phase, it is however apparent that regions with different LAT do indeed touch each other and therefore form a PD. Depending on the time resolution of the LAT map, staircase artifacts can be seen. However these can be filtered away by thresholding at an appropriately high value of *ρ*, as at those artifacts *ρ* is still much lower than at the PD.

To show the power of these methods, we applied them to a simulation and an OM experiment to find the length and orientation of PDLs over time. Here we conclude that using a method sensitive to PDLs allows to also identify linear rotor cores in experiment. However, having identified linear cores (PDLs) in optical voltage mapping of intact rabbit hearts [16] and human immortalized atrial myocyte cultures (in this work) does not allow to draw general conclusions. Therefore, we propose to use the suggested methods also on other datasets, starting with existing optical voltage mapping results. Then, the presented methods can be used to characterize PDL size and rotation. Such measurement would give another handle to judge the degree to which mathematical models of the heart resemble reality, in addition to e.g. reproducing restitution curves and observing basic spiral dynamics in terms of meander and stability.

We have verified that the proposed methods also work well when only noisy data or data at low resolution is available. In another experiment where the location of a thin, elongated obstacle was known as ground truth, we have seen that all proposed methods successfully are able to recover its location. We conjecture that therefore the methods also work well to recover PDLs in the setting of anatomical re-entry. More work is needed to better characterize the defect line analogues of functional and anatomical re-entry. In certain cases, the anchoring site may be a hybrid version of this classical distinction: Part of the linear rotor core can lie at an obstacle, or a functional region (PS or PDL) can be attracted to a inhomogeneity in the medium, to stay in place there, see Fig 16.

The methods used here are available as Python scripts from our online repositories, see the data availability statement. Please cite this paper when using the implementation. Note that some methods were originally introduced elsewhere: The AM method [23], the CM method [15] and RPG method [16]. The algorithms have currently been tested on dense Cartesian grids in 2D, but can naturally be extended to 3D and time, and unstructured grids (meshes), where the distinction between vertex-based densities *ρ* and edge-based densities *σ* will play a more prominent role.

## 6 Conclusion

In this work, we demonstrate that in order to visualize PDs in dense 2D data, it is recommended to use LAT-based methods or to use the skewed-phase to derive it from snapshots. Several algorithms were proposed to highlight the PDs visually, for which the simple methods (CM, RPG, CPG, PC) were most effective.

We applied the methods to simulations and an optical voltage mapping experiment; in the latter case we found that in a hiAM cell-culture, the average PDL length in a multi-spiral state was (4.01 *±* 0.55) mm and precession period *T* was (169.2 *±* 9.3) ms. We made our detection methods publicly available on our institutional repository and hope it can serve to further help understanding the building blocks of cardiac excitation patterns.

## Supporting information

In the supplementary material, we provide figures of PD densities *ρ, σ* for the different methods for AP and BOCF reaction kinetics, to enable a full comparison between methods.

The numerical methods implemented for this paper are available as a Python module at https://gitlab.com/heartkor/py_ithildin. The Python scripts used to generate the figures in this paper are available at https://gitlab.com/heartkor/scripts-pdl-detection. Finally, we have archived the simulation output and pre-processed optical voltage mapping data on which the scripts were applied on Zenodo (DOI: 10.5281/zenodo.6477532). This archive also contains the Python module and scripts.

Please cite this paper when using the implementation and/or the data.

**S1 Fig.**
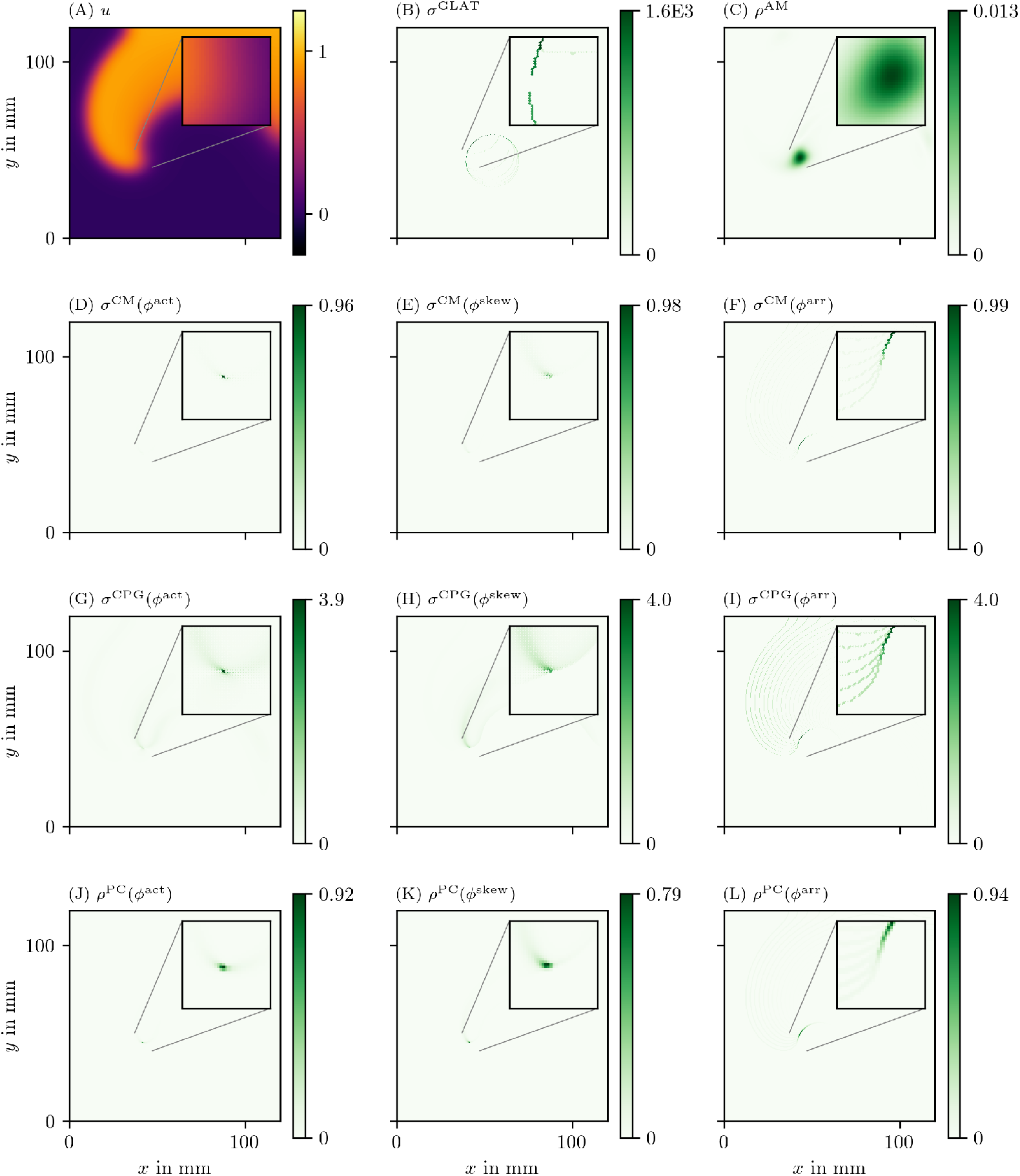
Overview of PD detection methods for one snapshot of the AP data set. The data are presented in the same way as in Fig 6.

**S2 Fig.**
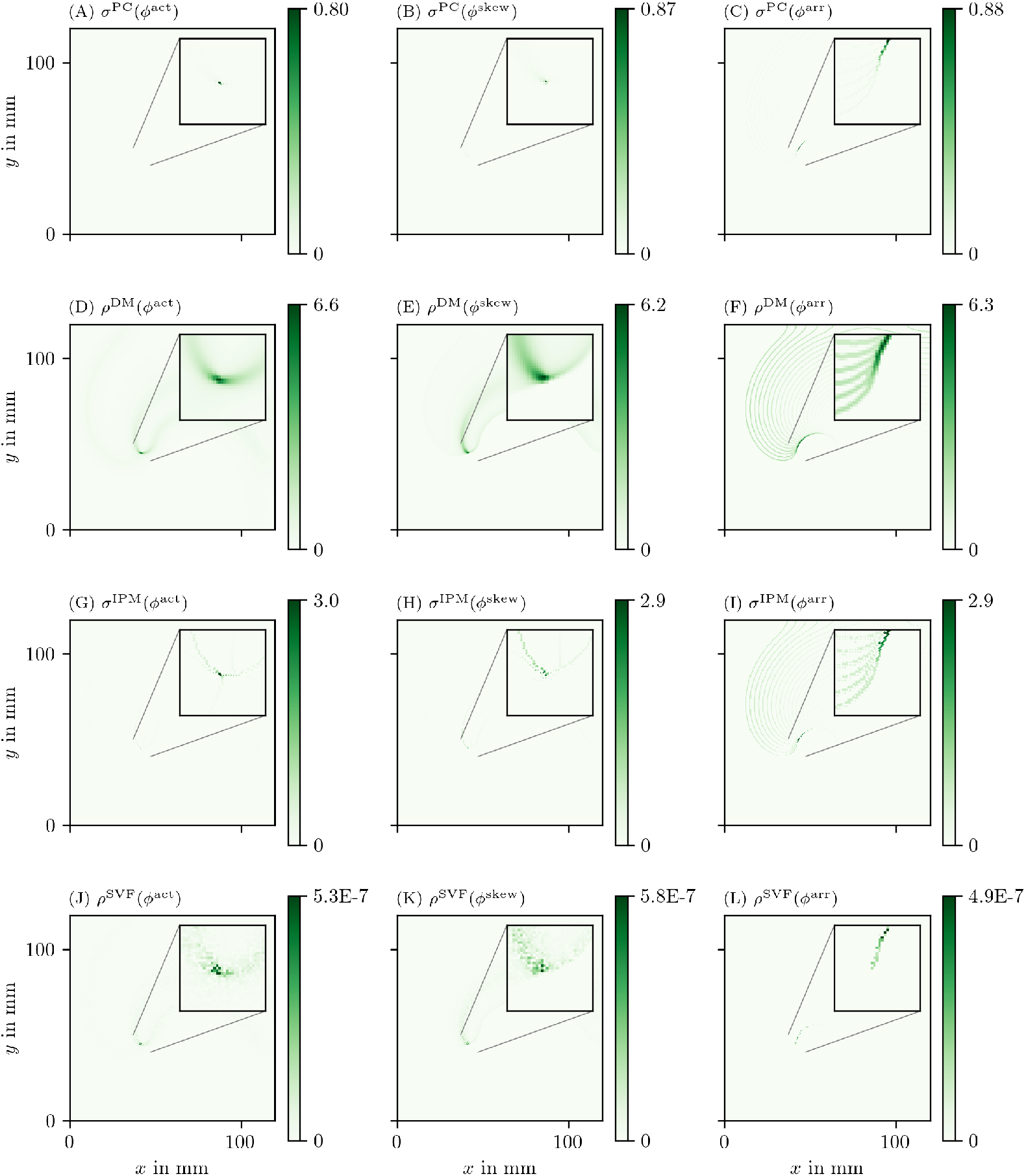
Overview of more PD detection methods for one snapshot of the AP data set as in S1 Fig.

**S3 Fig.**
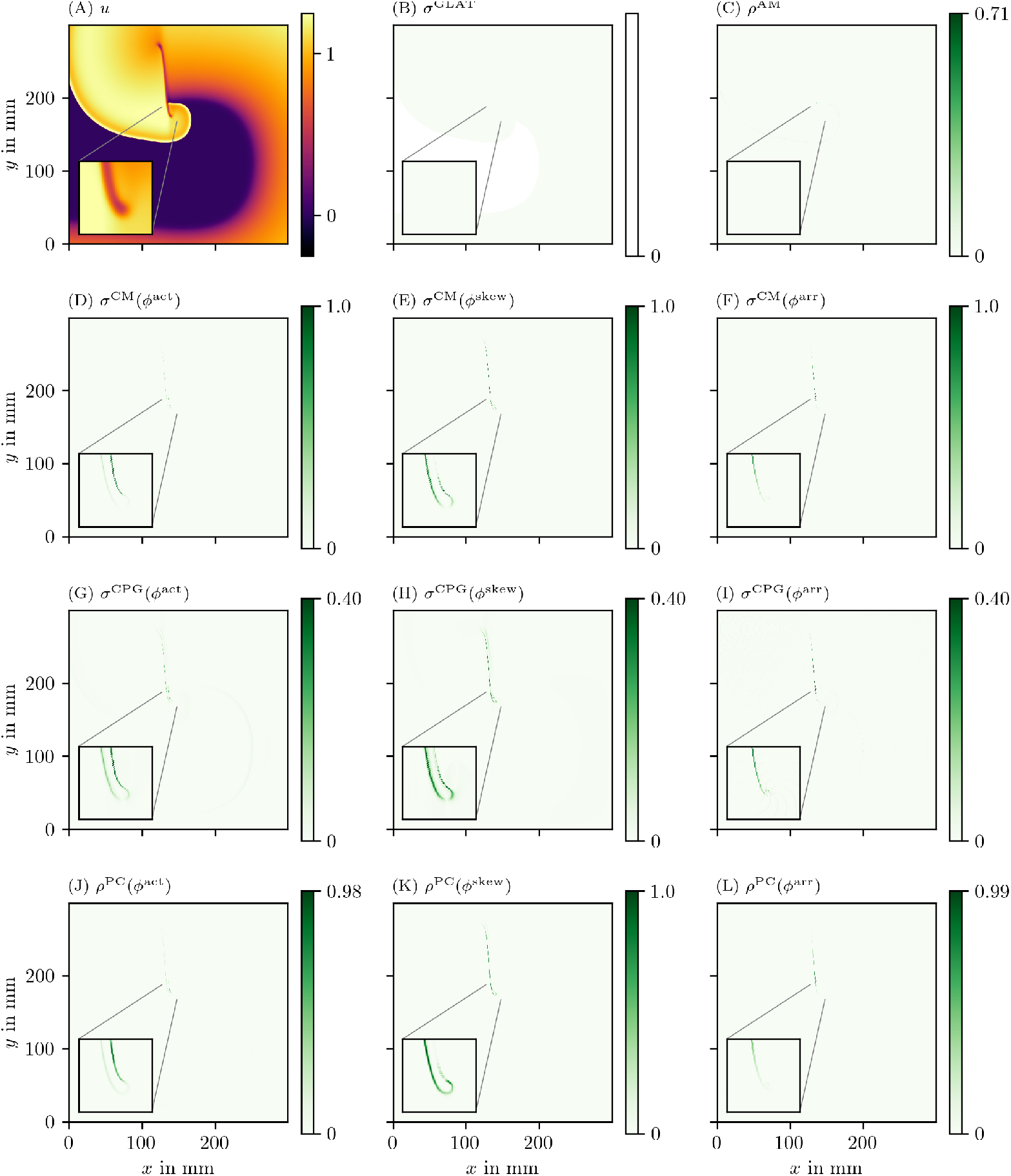
Overview of PD detection methods for one snapshot of the BOCF data set. The data are presented in the same way as in Fig 6.

**S4 Fig.**
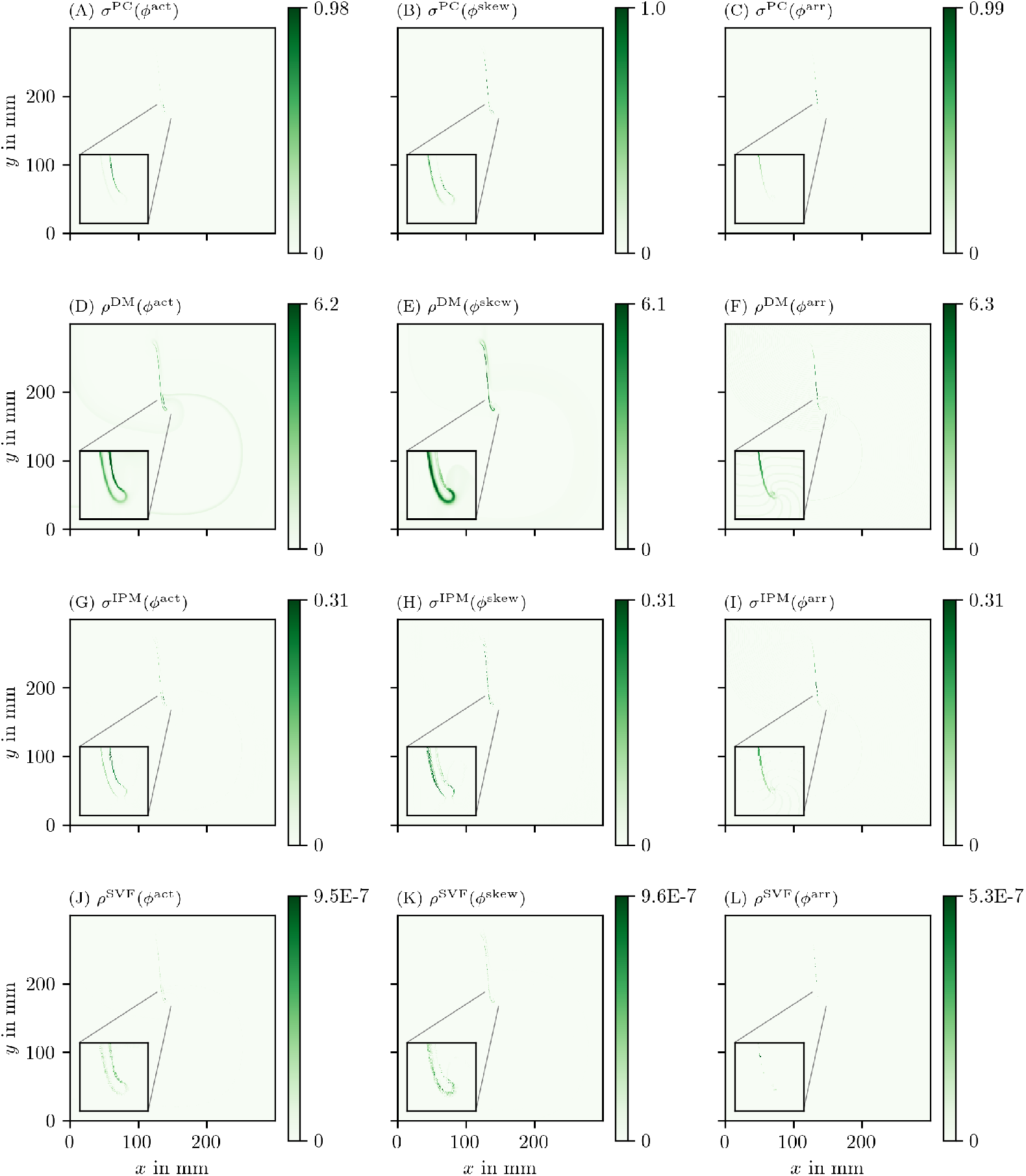
Overview of more PD detection methods for one snapshot of the BOCF data set as in S3 Fig.

## Acknowledgments

We are grateful to Sven O. Dekker, Niels Harlaar, Daniël A. Pijnappels and Antoine A.F. de Vries for providing optical voltage mapping data of cardiomyogenically differentiated hiAM monolayers. Moreover, we thank Tim De Coster for helpful comments on the analogy between a PDL and the spiral wave tip trajectory.

## Author contribution

HD and AVP conceived the study. DK, LA and LL ran the numerical simulations and implemented the detection algorithms. All authors wrote parts of the manuscript and assisted in internal reviewing.

## Funding

DK is supported by KU Leuven grant GPUL/20/012. LA was funded by a KU Leuven FLOF grant and a FWO-Flanders fellowship, grant 1177022N; LL was funded by KU Leuven and FWO-Flanders, grant G025820N. Research at Sechenov University was financed by the Ministry of Science and Higher Education of the Russian Federation within the framework of state support for the creation and development of World-Class Research Centers “Digital biodesign and personalized healthcare” 075-15-2020-926.

## Notes

https://gitlab.com/heartkor/py_ithildin

https://gitlab.com/heartkor/scripts-pdl-detection

https://doi.org/10.5281/zenodo.6477532

### Competing Interest Statement

The authors have declared no competing interest.

### Summary of Updates

Updated preprint for re-submission to PLOS One. Reviewer comments have been applied and new in silico experiments conducted investigating noise, low spatial resolution, and an experiment with a known ground truth.

